# Non-canonical internalization mechanisms of mGlu receptors

**DOI:** 10.1101/2025.02.13.638043

**Authors:** M Cimadevila, J Liu, D Maurel, I Brabet, M Hoscar, J Drube, C Hoffmann, A Inoue, P Rondard, PA Lafon, L Prézeau, JP Pin

## Abstract

Cell surface density of G protein-coupled receptors (GPCRs) is tightly regulated through constitutive and agonist-induced internalization. Whereas the mechanisms of constitutive internalization remain elusive, agonist-induced internalization is accepted to involve receptor phosphorylation by GPCR kinases (GRKs), β-arrestin binding and AP2 recruitment, targeting receptors to clathrin-coated pits. Dimeric class C metabotropic glutamate (mGlu1 to 8) receptors regulate synaptic transmission but their internalization process is ambiguous. Here, we used diffusion-enhanced energy transfer (DERET) to decipher their internalization kinetics. We showed that all mGlu receptors are constitutively internalized. However, only mGlu1, 5 and 3 homodimers are agonist-induced internalized, that require neither GRKs, nor β-arrestins. In contrast, the constitutive internalization involves only β-arrestins. This systematic study further illustrates how different class C receptors are relative to most other GPCRs, revealing non-canonical internalization mechanisms. These insights in mGlu receptor dynamics will help promoting the therapeutic action of drugs targeting mGlu receptors.

## Introduction

G protein-coupled receptors (GPCRs) are seven transmembrane domain proteins that constitute the target of around 30 % of the drugs on the market^1^. They are mainly embedded within the lipid bilayer of the cell membrane but as they are highly dynamic, they can internalize to intracellular compartments. This process can happen in the absence of ligands (constitutive internalization) or upon agonist binding (agonist-induced internalization) through potentially different mechanisms.

Agonist-induced internalization is canonically described as a clathrin-mediated process^2^, although other mechanisms have been described including caveolae-mediated ones^3^. Binding of an agonist leads to GPCR phosphorylation by GPCR kinases (GRKs) and β-arrestin (βarr) recruitment. As a consequence, βarrs are activated and trapped as cargos by the protein adaptor complex 2 (AP2). AP2 drives the invagination of the activated GPCR in clathrin-coated pits^4^. In comparison, the mechanisms of GPCR constitutive internalization remain elusive, being mostly described as non-clathrin mediated^3,5,6^.

For class C GPCRs such as metabotropic glutamate receptors (mGlu receptors; mGlu1 to mGlu8), multiple studies report on their desensitization mechanisms^7,8^. Yet, their internalization is still largely unknown and controversial. While it has recently been reported that some mGlu receptors can internalize following the canonical internalization pathway^9–11^, other reports suggested that mGlu receptors can be internalized by a clathrin-independent pathway^12–14^. This knowledge is key due to the fact that mGlu receptors are widely studied as targets for modulating chronic brain diseases, such as Parkinson disease, schizophrenia or Alzheimer’s disease^15^. However, drug candidates have failed in clinical trials mainly due to their lack of efficacity^16^. As long-term treatments rely on the continuous presence of the target, internalization might compromise chronic effect of drugs activating mGlu receptors.

This is why we aimed at precisely determining the internalization profiles and mechanisms of mGlu receptor internalization. To that end, we used diffusion-enhanced resonance energy transfer (DERET) combined with protein knock-out or down-regulation. We showed that mGlu receptors internalize through a new non-canonical mechanism, although it involves the classical partners. Upon agonist-binding, mGlu1, 5, and 3 internalize in a βarr-independent manner that absolutely requires AP2. In contrast, constitutive internalization is dependent on βarr and precisely shaped by GRKs. This information provides fundamental insights for the analysis of the possible impact of their internalization on therapeutic strategies and ways to regulate it.

## Results

### Identification of mGlu5 internalization kinetics

Internalization kinetics have been measured by DERET, an approach that takes advantage of the quenching of the long-life emission of lanthanides by diffusible acceptors at high concentration. To measure internalization by DERET, GPCRs at the plasma membrane are covalently labelled with a lanthanide through a suicide enzyme genetically introduced at their N-termini. The addition of a saturating concentration of fluorescein, the acceptor, quenches the fluorescence of the lanthanide^17,18^. If the GPCR is internalized, the lanthanide is physically separated from the acceptor and so its fluorescence is recovered. Fluorescence recovery can then be recorded over time to identify the GPCR internalization kinetics (Fig. 1A).

**Figure 1.**
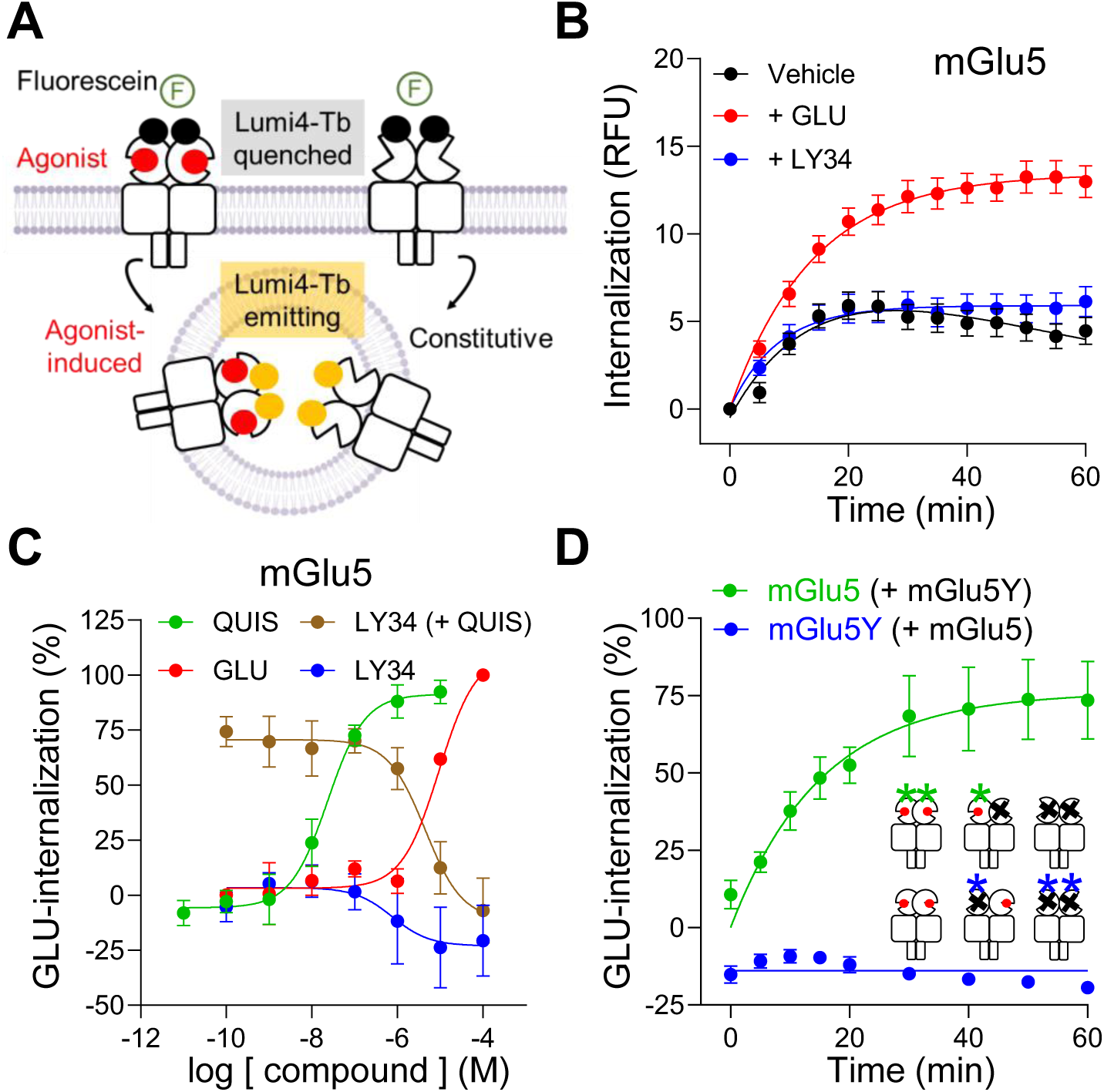
DERET revealed internalization properties of mGlu5. (**A**) Schema of the experimental approach used to measure internalization kinetics by DERET. (**B**) Internalization kinetics of mGlu5 in the absence of agonist (Vehicle), in the presence of 100 μM glutamate (+ GLU) or in the presence of 100 μM LY341495 (+ LY34). (**C**) Agonist-induced internalization measured after 60 min in the presence of increasing doses of glutamate (+ GLU), quisqualate (+ QUIS), LY341495 (+ LY34) or LY341495 (+ 100 nM quisqualate). Data were normalized to glutamate-induced internalization after 60 min of incubation at 37 °C. (**D**) Agonist-induced internalization (1 μM quisqualate) of mGlu5 when co-expressed with mGlu5Y (green line line) and mGlu5Y when co-expressed with mGlu5 (blue line). Data were normalized to glutamate-induced internalization of mGlu5-only expressing cells. Data are means ± S.E.M. of at least three independent experiments performed in triplicates.

Accordingly, mGlu5 was labelled thanks to a SNAP-suicide enzyme domain inserted at its N-terminus after the signal peptide. Addition of a SNAP-substrate linked to Lumi4-terbium (Lumi4-Tb) allowed the covalent labelling at 4 °C of the receptors present at the plasma membrane (Fig. S1). Internalization was then started by the addition of the indicated ligands and fluorescein and followed for 60 min at 37 °C. Data were corrected by receptor expression as explained in material and methods, and fitted to previously described pharmacokinetic equations^19^.

mGlu5 was constitutively internalized as an increase in Lumi4-Tb luminescence was observed in the absence of agonist and the presence of an antagonist (Fig. 1B). Such an effect is not due to the possible action of ambient glutamate in the media up to 45 min, as internalization is not inhibited by the competitive antagonist LY341495 (Fig. 1B). The endogenous agonist glutamate was found to increase internalization (Fig. 1B), in a concentration-dependent manner (Fig. 1C), same as for the selective agonist quisqualate (Fig. 1C), with EC_50_ values similar to the ones found in the literature^20^ (Table S1). Also, the antagonist LY341495 is able to block the agonist-induced internalization induced by quisqualate with the expected potency (Fig. 1C, table S1). Note that LY341495 has the tendency to decrease the basal mGlu5 internalization after 60 min of incubation at 37 °C, an effect that may likely be due to variable levels of ambient glutamate accumulation in the assay media^21^. Accordingly, the basal internalization will always be measured in the presence of an antagonist to prevent any possible effect of ambient glutamate.

As mGlu receptors are mandatory dimers, two identical (homodimers) or different (heterodimers) subunits can be involved in agonist-induced internalization^22^. We wondered whether both subunits of the dimers should be in an active state for the receptor to undergo agonist-induced internalization. To investigate this, we co-expressed mGlu5 with a mutated subunit that is unable to bind glutamate, mGlu5Y^23^, to examine the internalization profile of the heterodimer mGlu5-5Y. Three populations of receptors were expected: mGlu5-5, mGlu5Y-5Y, and mGlu5-5Y. By the use of two different suicide enzymes to tag mGlu5 and mGlu5Y, we could selectively follow internalization of one subunit or the other, and so evaluate the internalization properties of the dimer. We found that mGlu5Y was unable to be agonist-induced internalized when co-expressed with mGlu5 (Fig. 1D), even though they can efficiently dimerize (Fig. S2). Agonist-induced internalization of mGlu5 requires then to have both subunits in the active conformation.

### Only three out of eight homodimers are agonist-induced internalized

The 8 mGlu subtypes are classified into three groups according to their sequence homology. Group-I includes subtypes 1 and 5; group-II includes subtypes 2 and 3; group-III includes subtypes 4, 6, 7 and 8. Once DERET was validated as a valuable tool for monitoring the internalization of mGlu receptor dimers, we explored the internalization profile of all mGlu receptors. Homodimers internalization was investigated by expressing the SNAP-tagged mGlu subtypes in HEK293 cells (Fig. S3). All homodimers were constitutively internalized, as in the absence of agonist they show an increase of internalization over time (Fig. 2, Basal). Constitutive internalization increases over time except for mGlu5 and mGlu3, whose constitutive internalization reaches a plateau after 20 min.

**Figure 2.**
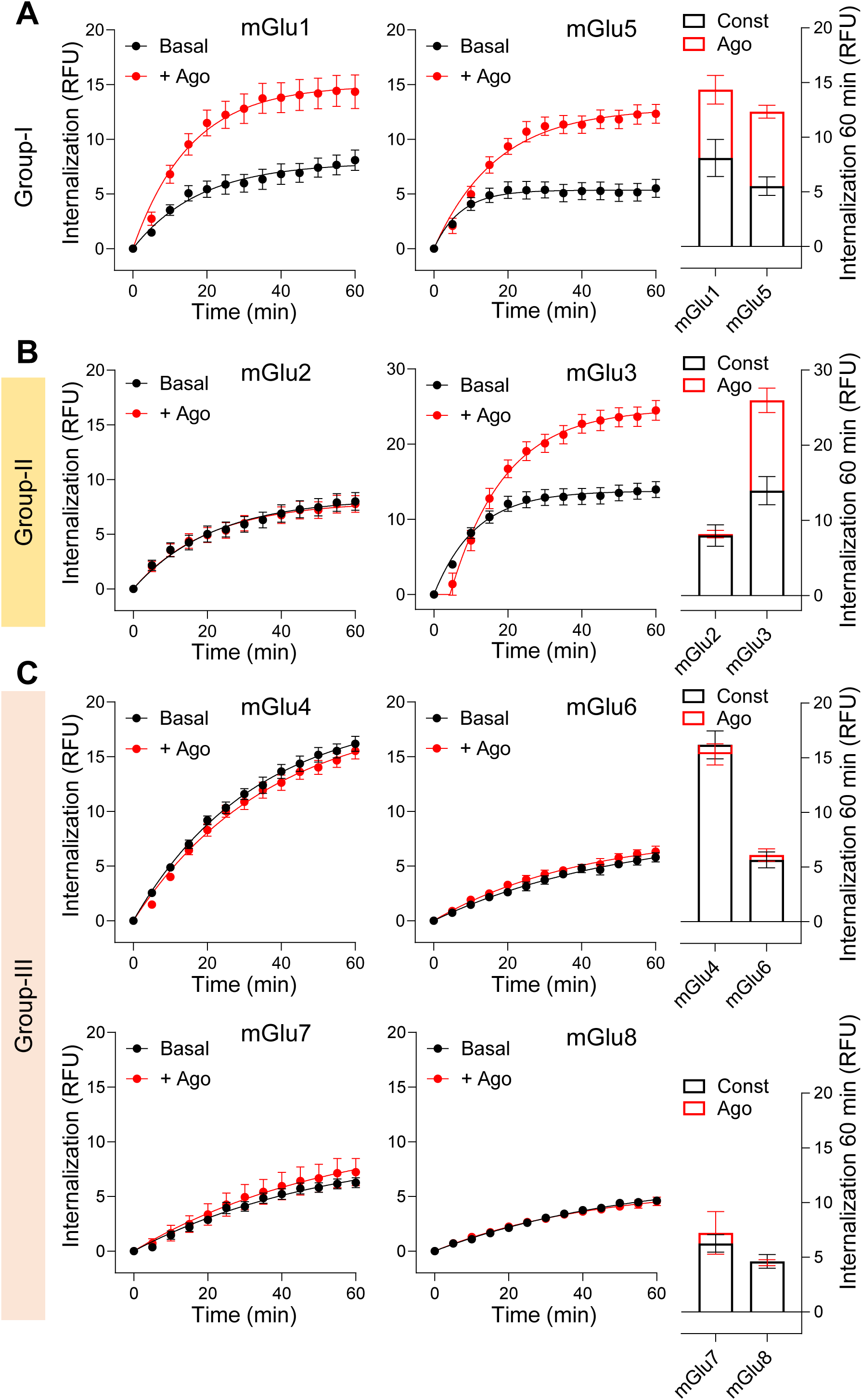
Kinetic analysis reveals different internalization profiles of mGlu homodimers. (**A**) Internalization profiles of group-I mGlu receptors, mGlu1 and mGlu5 in the absence of agonist (basal, 100 μM LY341495) or 10 μM quisqualate (+ Ago). (**B**) Internalization profile of group-II mGlu receptors, mGlu2 and mGlu3, in the absence of agonist (basal, 100 μM LY341495) or 10 μM LY354740 (+ Ago). Bars represent constitutive internalization (const, given by the basal signal) and agonist induced (ago, calculated by subtracting basal signal) after 60 min of internalization at 37 °C. (**C**) Internalization profile of group-III mGlu receptors, mGlu4, mGlu6, mGlu7, mGlu8, in the absence of agonist (basal, 100 μM LY341495) or 10 μM L-AP4 for mGlu4, mGlu6, mGlu8; 10 μM LSP2-9166 for mGlu7 (+ Ago). Bars represent constitutive internalization (const, given by the basal signal) and agonist induced (ago, calculated by subtracting basal signal) after 60 min of internalization at 37 °C. Data represent the means ± S.E.M. of at least three independent experiments performed in triplicates.

However, none of the group-III receptor basal internalization was increased with a saturating dose of agonist (Fig. 2C; and Fig. S4). Specifically, among the eight mGlu homodimers, only three showed agonist-induced internalization: both group-I mGlu1 and 5, and the group-II mGlu3 receptors (Fig. 2; Fig. S4). It was surprising to see that upon agonist activation, no mGlu3 internalization could be detected within the first 5 min, meaning that even the constitutive internalization is blunted (Fig. 2B). This delay was systematically observed (see Fig. 5C, 6C, and 8B). Certainly, addition of an agonist increased internalization over basal until the equilibrium, which is reached at around 30 min of agonist incubation. Compensatory mechanisms, such as recycling, could explain these kinetic profiles.

### Agonist-induced internalized receptor undergoes recycling

Recycling refers to the process where a GPCR, which has been located in the plasma membrane and then internalized, comes back to the plasma membrane^24^. To evaluate mGlu receptor recycling, labelled mGlu5 and mGlu3 were incubated with an agonist during 60 min at 37 °C to induce their internalization. Then, either the excess of agonist was removed or the antagonist LY341495 was added, as antagonists have been reported to accelerate recycling^25^. If the receptor is recycled to the plasma membrane, Lumi-4-Tb will be quenched at the surface (Fig. 3A).

**Figure 3.**
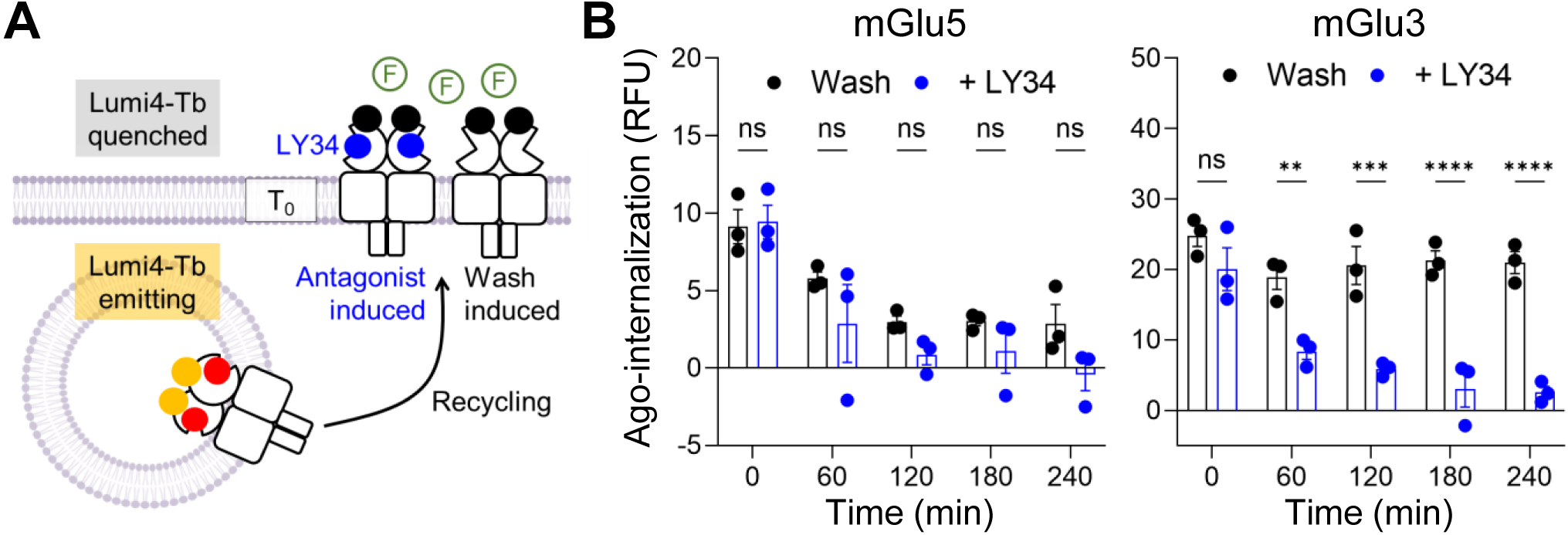
Different recycling properties of internalized mGlu3 and mGlu5 receptors. (**A**) Schema of the experimental approach used to measure recycling of agonist-induced internalized receptors. Receptor internalization was induced for 60 min at 37 °C. At time 0 (T_0_), the excess of agonist was removed, and either vehicle (wash) or LY34 were added in the presence of fluorescein. Internalization was measured every 60 min from T_0_ at 37 °C. (**B**) Remaining intracellular agonist-induced internalized mGlu5 (left) or mGlu3 (right) (10 μM QUIS, or 10 μM LY354740, respectively) with time after washing the agonist (wash) or adding 100 μM LY341495 (+ LY34). Data represent the means ± S.E.M. of at least three independent experiments performed in triplicates.

Indeed, mGlu5 was recycled back to the plasma membrane over time, and the addition of LY341495 did not significantly increase recycling when compared with only washing the agonist (Fig. 3B). Similarly, mGlu3 recycling was observed upon LY341495 antagonist application (Fig. 3B), in contrast to the washing condition alone. We cannot exclude the possibility that, after washing, ambient glutamate is accumulating over time and reaches the concentration to activate mGlu3, due to the high affinity of this transmitter for mGlu3^21^. The identification of receptor recycling after agonist induced internalization highlights the relevance of measuring kinetics while evaluating mGlu receptor internalization.

### Only mGlu2-3 among the mGlu3-containing heterodimers internalizes upon agonist stimulation

It is now well accepted that mGlu receptors exist not only as homodimers, but also as heterodimers with specific subunit composition^22,26^. While group-I mGlu receptors only form heterodimers between themselves, both group-II and group-III can associate into any possible dimer combination. Among them, only mGlu3 homodimer was found to undergo agonist-induced internalization. We then examined whether mGlu3-containing heterodimers could be agonist-induced internalized.

By co-expressing SNAP-mGlu3, with CLIP-mGlu2, CLIP-mGlu4, CLIP-mGlu7 or CLIP-mGlu8, we were able to only label the non-agonist-induced internalizing subunit of the heterodimer (mGlu2, mGlu4, mGlu7 and mGlu8) and follow its internalization over time (Fig. 4A). We verified that heterodimers could be detected at the cell surface, at a different extent, using time-resolved fluorescence resonance energy transfer (TR-FRET) compatible substrates for SNAP and CLIP (Fig. 4B). Intriguingly, only mGlu2 subunit is agonist-induced internalized in the presence of mGlu3 (Fig. 4C), but not mGlu4, mGlu7 or mGlu8 (Fig. 4D). This indicates that mGlu2-3 is the only mGlu3-containing heterodimer to be agonist-induced internalized, adding this receptor to the mGlu entities that are internalized upon activation.

**Figure 4.**
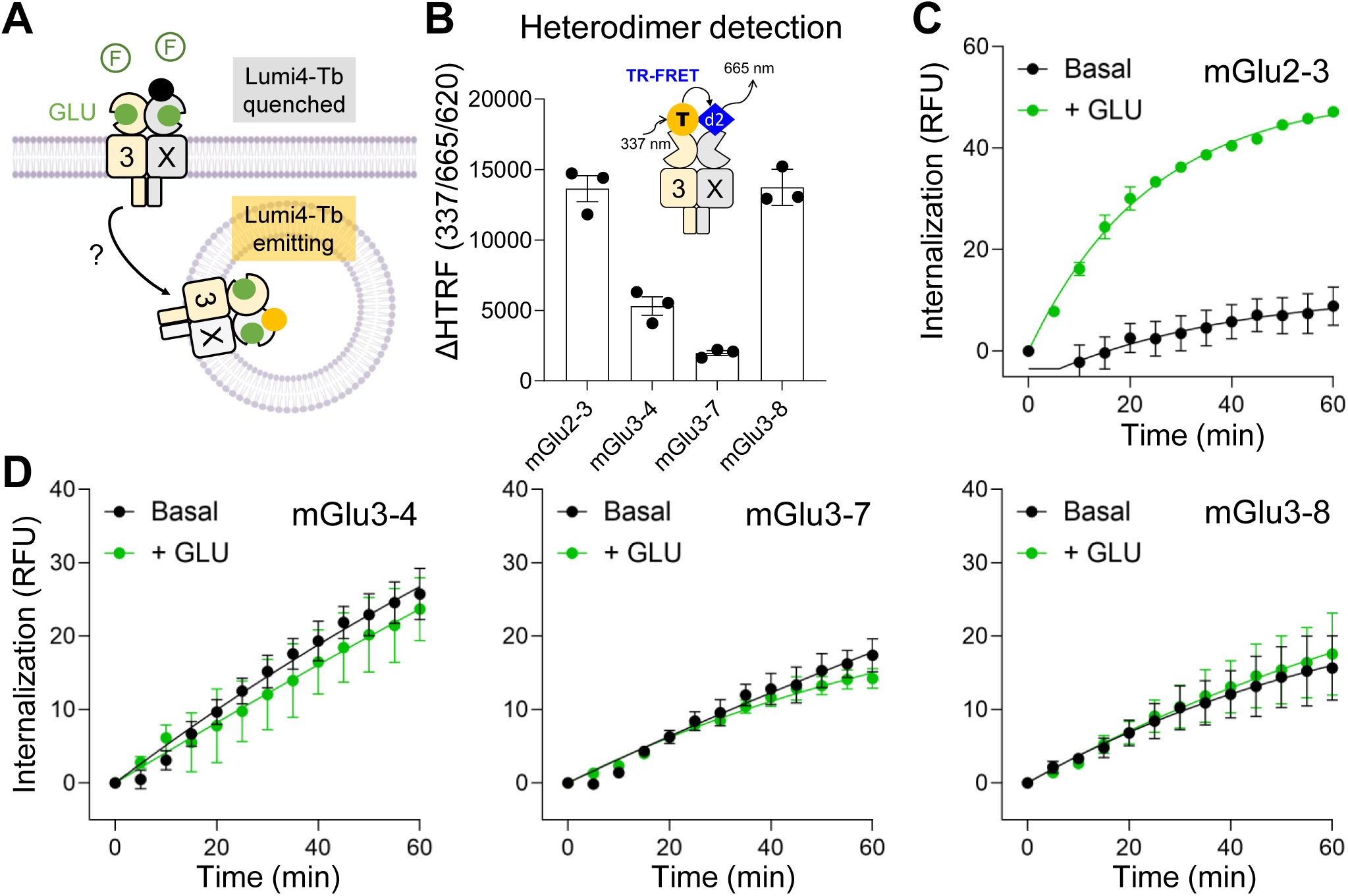
Only mGlu2-3 among the mGlu3-containing heterodimers internalizes upon agonist stimulation. (**A**) Schema of the experimental approach used to measure heterodimer internalization by selectively tag one of the two subunits of the dimer. (**B**) TR-FRET between mGlu3 subunit labelled with Lumi4-Tb (T) and mGlu2, mGlu4, mGlu7 or mGlu8 labelled with d2. ΔHTRF™ (337/665/620) represents the specific TR-FRET signal calculated by subtracting TR-FRET signal obtained in non-transfected cells. (**C**) Internalization of mGlu2 when co-expressed with mGlu3, in the absence of agonist (basal, 100 μM LY341495) or in the presence of 100 μM glutamate (+ GLU). (**D**) Internalization of group-III mGlu receptors, mGlu4, mGlu7 or mGlu8 when co-expressed with mGlu3, in the absence of agonist (basal, 100 μM LY341495) or in the presence of 100 μM glutamate (+ GLU). Data represent the means ± S.E.M. of at least three independent experiments performed in triplicates.

### GRKs are not mandatory for agonist-induced internalization

Three main molecular players are involved in the canonical GPCR internalization process: GRKs, βarr and AP2. GRKs are kinases that can be dispersed in the cytoplasm (GRK2 and GRK3) or associated with the plasma membrane (GRK5 and GRK6). They phosphorylate the C-terminal domain of agonist-bound GPCRs as a first step in the internalization process for most GPCRs (Fig. 5A), such as μ-opioid receptors (MOR)^27^. Certainly, inhibition of GRK2/3 with the GRK inhibitor cmpd101 decreased agonist-induced internalization of MOR without affecting constitutive internalization (Fig. 5B). The complete deletion of GRKs (ΔQ-GRK cell line^28^, abbreviated as ΔGRK) blunted agonist-induced internalization of MOR, internalization that could be rescued by overexpression of any of the GRKs (Fig. 5B). In contrast, MOR constitutive internalization was independent of GRKs (Fig. 5B).

**Figure 5.**
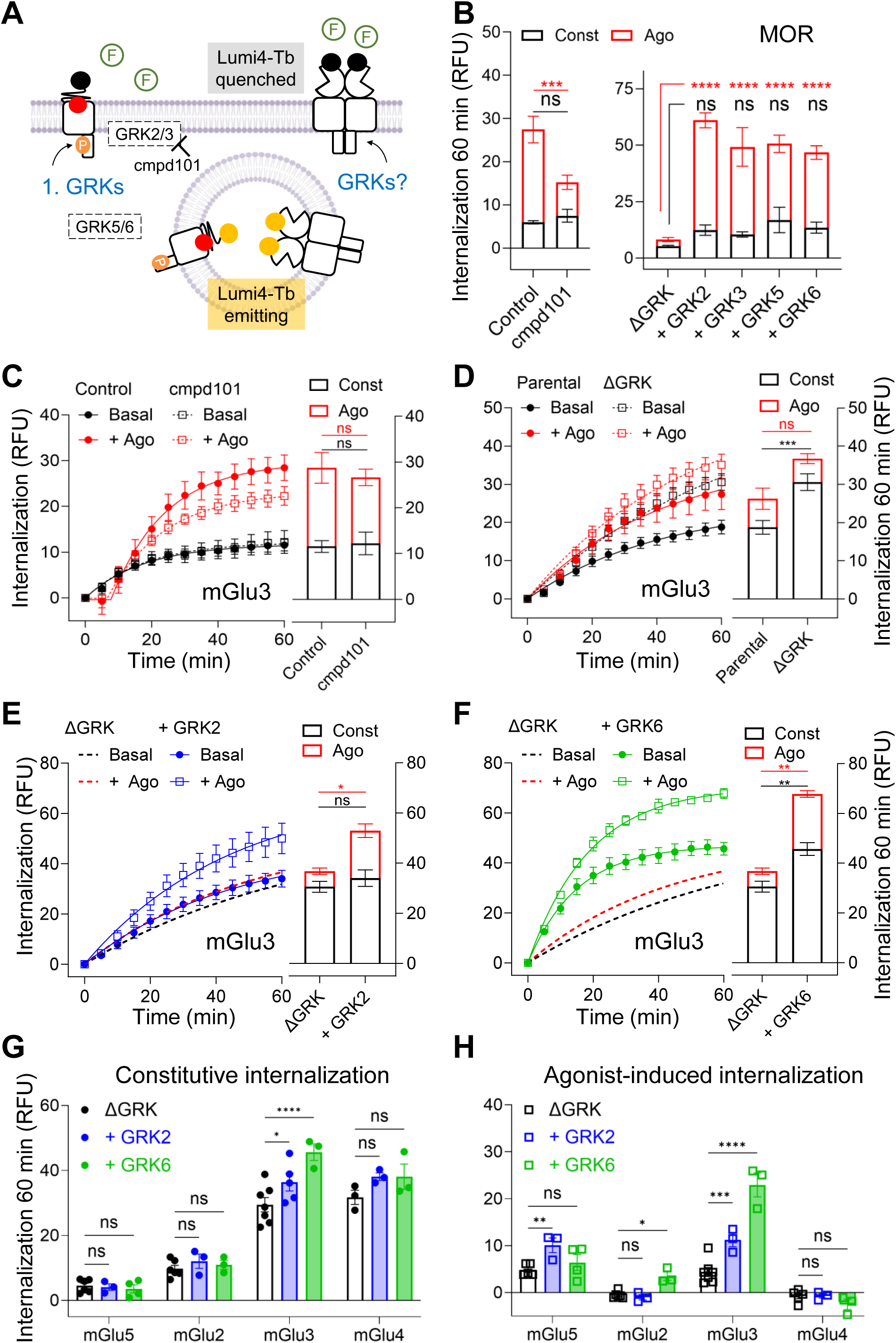
GRKs are not mandatory for mGlu receptor internalization. (**A**) Schema representing canonical impact of GRKs in GPCR internalization. (**B**) MOR internalization measured after 60 min in the absence of agonist (constitutive) or in the presence of 37.5 μM of DAMGO (agonist-induced; ago). Experiments were performed in HEK293 cells (control), in HEK293 cells pre-treated with 10 μM of cmpd101 during 30 min at 4 °C; in cells lacking of all GRKs under control condition (ΔGRK) or supplemented with a single GRK (+ GRK2, + GRK3, + GRK5, + GRK6). (**C**) Internalization of mGlu3 receptor in the absence (basal, 100 μM LY341495) or in the presence of agonist (+ Ago, 10 μM LY354740), without or with a pretreatment of 10 μM of cmpd101 during 30 min at 4 °C. Bars represent constitutive internalization (const, given by the basal signal) and agonist induced (ago, calculated by subtracting basal signal) after 60 min of internalization at 37 °C. (**D**) Internalization of mGlu3 receptor in the absence (basal, 100 μM LY341495) or in the presence of agonist (+ Ago, 10 μM LY354740), in parental or ΔGRK cells. Bars represent constitutive internalization (const, given by the basal signal) and agonist induced (ago, calculated by subtracting basal signal) after 60 min of internalization at 37 °C. Internalization of mGlu3 in the absence (basal, 100 μM LY341495) or in the presence of agonist (+ Ago, 10 μM LY354740), in ΔGRK cells + GRK2 (**E**), + GRK6 (**F**). Bars represent constitutive internalization (const, given by the basal signal) and agonist induced (ago, calculated by subtracting basal signal) after 60 min of internalization at 37 °C. (**G**) Constitutive internalization of mGlu5, mGlu2, mGlu3 and mGlu4 after 60 min treatment with 100 μM LY341495, in ΔGRK cells (black), in ΔGRK cells + GRK2 (blue) or in ΔGRK cells + GRK6 (green). (**H**) Agonist-induced internalization of mGlu5, mGlu2, mGlu3 and mGlu4 after 60 min treatment with a selective agonist (10 μM quisqualate for mGlu5, 10 μM LY354740 for mGlu2 and mGlu3, 10 μM LAP4 for mGlu4), in ΔGRK cells (black), in ΔGRK cells + GRK2 (blue) or in ΔGRK cells + GRK6 (green). Data represent the means ± S.E.M. of at least three independent experiments performed in triplicates.

We then unveiled the role of GRKs in both strong constitutive and agonist-induced internalization of mGlu3. Cmpd101 failed to induce a change of mGlu3 internalization, both constitutive and agonist-induced ones (Fig. 5C). Also, in the absence of GRKs, agonist-induced internalization of mGlu3 was preserved and constitutive internalization was even increased (Fig. 5D). To assess if GRKs individually could impact internalization of mGlu3, we overexpressed GRK2 or GRK6, as they are highly expressed in HEK293 cells^29^ and in the human brain^30^. GRK2 boosted agonist-induced internalization (Fig. 5E), while GRK6 boosted both constitutive and agonist-induced internalization (Fig. 5F). GRK effect was kinase dependent as kinase dead mutants had no effect (Fig. S5).

As the two different GRKs had a different impact in mGlu3 internalization, we extended the study to mGlu5 as example of agonist-induced group-I mGlu receptors; to mGlu2, a group-II receptor with different internalization properties than mGlu3, and to mGlu4 as an example of group-III mGlu receptors. We compared internalization in the absence (ΔGRK) or presence of GRK2 or GRK6 after 60 min. In contraposition of mGlu3, neither mGlu5, mGlu2 or mGlu4 constitutive internalization was affected by GRKs (Fig. 5G). Regarding to agonist-induced internalization, mGlu2 kept GRK-preference for GRK6 over GRK2, as significant agonist-induced internalization was revealed in the presence of GRK6 (Fig. 5H). Intriguingly, mGlu5 agonist-induced internalization was only boosted by GRK2 and not GRK6 (Fig. 5H). Similar data was obtained with GRK3 and GRK5 (Fig. S6). Taken together, although dispensable for mGlu receptor internalization, GRKs can tune internalization of the studied mGlu receptors.

### β-arrestins are only required for constitutive internalization

βarr1 and βarr2 bind to GRK-phosphorylated GPCRs to start their internalization (Fig. 6A). In accordance, removal of βarr (Arr-2/3-null cell line^31^ abbreviated as Δβarr) blunted agonist-induced internalization of vasopressin receptor 1B (V_1b_). Internalization was recovered after overexpression of βarr1 or βarr2 (Fig. 6B). Constitutive internalization happens independently of βarrs for V1b receptor as expected (Fig. 6B).

**Figure 6.**
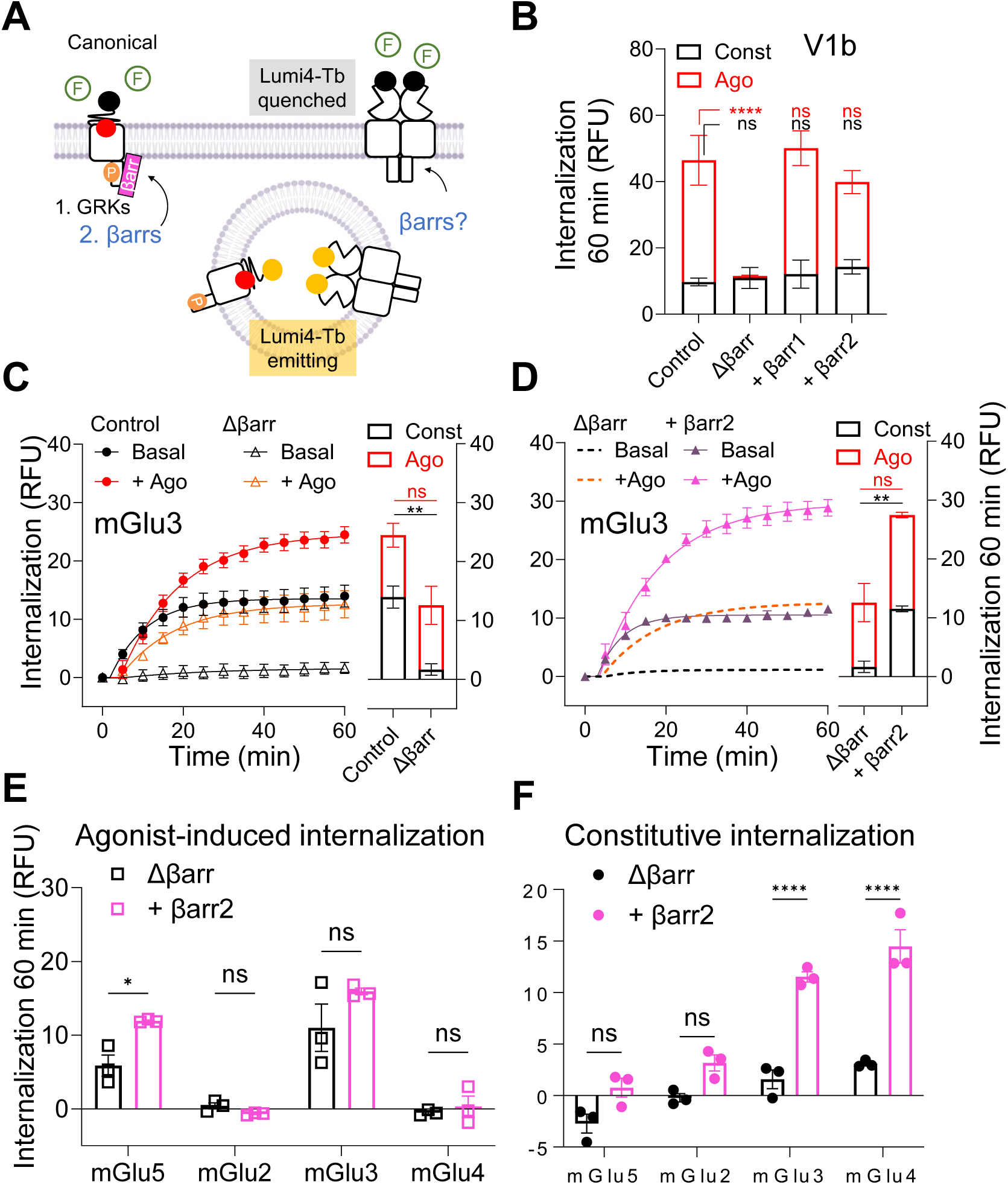
βarrs are mandatory for constitutive mGlu receptor internalization only. (**A**) Schema representing canonical impact of βarrs in GPCR internalization. (**B**) V_1b_ internalization measured after 60 min in the absence of agonist (constitutive) or in the presence of 10 μM of SR149515 (agonist-induced; ago) in HEK293 cells (control), Δβarr cells, Δβarr cells and Δβarr cells supplemented with + βarr1 (+ βarr1) or βarr2 (+ βarr2). (**C**) Internalization kinetics of mGlu3 receptor in HEK293 cells (control) and in Δβarr cells in the presence of 100 μM LY341495 (basal) or 10 μM LY354740 (+ Ago). Bars represent constitutive internalization (const, given by the basal signal) and agonist induced (ago, calculated by subtracting basal signal) after 60 min of internalization at 37 °C. (**D**) Internalization kinetics of mGlu3 receptor in Δβarr cells supplemented with βarr2 in the presence of 100 μM LY341495 (basal) or 10 μM LY354740 (+ Ago). Bars represent constitutive internalization (const, given by the basal signal) and agonist induced (ago, calculated by subtracting basal signal) after 60 min of internalization at 37 °C. (**E**) Agonist-induced internalization of mGlu5 (+ 10 μM quisqualate), mGlu2 (+ 10 μM LY354740), mGlu3 (+ 10 μM LY354740) or mGlu4 (+ 10 μM L-AP4) after 60 min in Δβarr cells (black bars) and in Δβarr cells supplemented with βarr2 (pink bars). (**F**) Constitutive internalization of mGlu5, mGlu2, mGlu3 and mGlu4 measured after 60 min of incubation with in 100 μM LY341495 in Δβarr cells (black bars) and in Δβarr cells supplemented with βarr2 (pink bars). Data represent the mean ± S.E.M. of at least three independent experiments performed in triplicate.

As mGlu receptors are internalized independently of GRKs, we expected no impact of βarrs on their internalization. Indeed, agonist-induced internalization of mGlu3 (Fig. 6C) and mGlu5 (Fig. 6F) was observed in Δβarr cells (Fig. 6), despite their correct expression (Fig. S7). Still, overexpression of either βarr1 or βarr2 in these cells increased agonist-induced internalization of mGlu5 (Fig. 6E; Fig. S8-S9) and of mGlu3 when overexpressed with βarr1 (Fig. S9). Surprisingly constitutive internalization of all mGlu receptors tested (mGlu2, 3, 4 and 5) was largely decreased in the Δβarr cells (Fig. 6C,F), but restored when βarr1 or βarr2 were added (Fig. 6D; Fig. S8-S9).

To further investigate this effect, we evaluated the recruitment of βarrs by the receptors through bioluminescence resonance energy transfer (BRET). We tagged the C-terminal domain of δ-opioid receptor (DOR) used as a prototypical class A GPCR, and mGlu5, mGlu2, mGlu3 and mGlu4 with the fluorescent protein Venus (Fig. 7A). By co-expressing each construct with βarr1 or βarr2 tagged with luciferase (RLuc), we were able to follow βarr recruitment by the receptors over time. Addition of a selective agonist only increased recruitment of βarr2 by DOR, without having any effect on mGlu receptors (Fig. 7B). Interestingly, a large basal BRET was measured between mGlu3 and βarr2 similar to that observed with DOR after agonist activation, an effect that is not affected upon mGlu3 activation (Fig. 7B), nor by the antagonist LY341495 (Fig. S10A-B) indicating this effect is not due to ambient glutamate. This effect was also observed with βarr1 (Fig. S10C-E).

**Figure 7.**
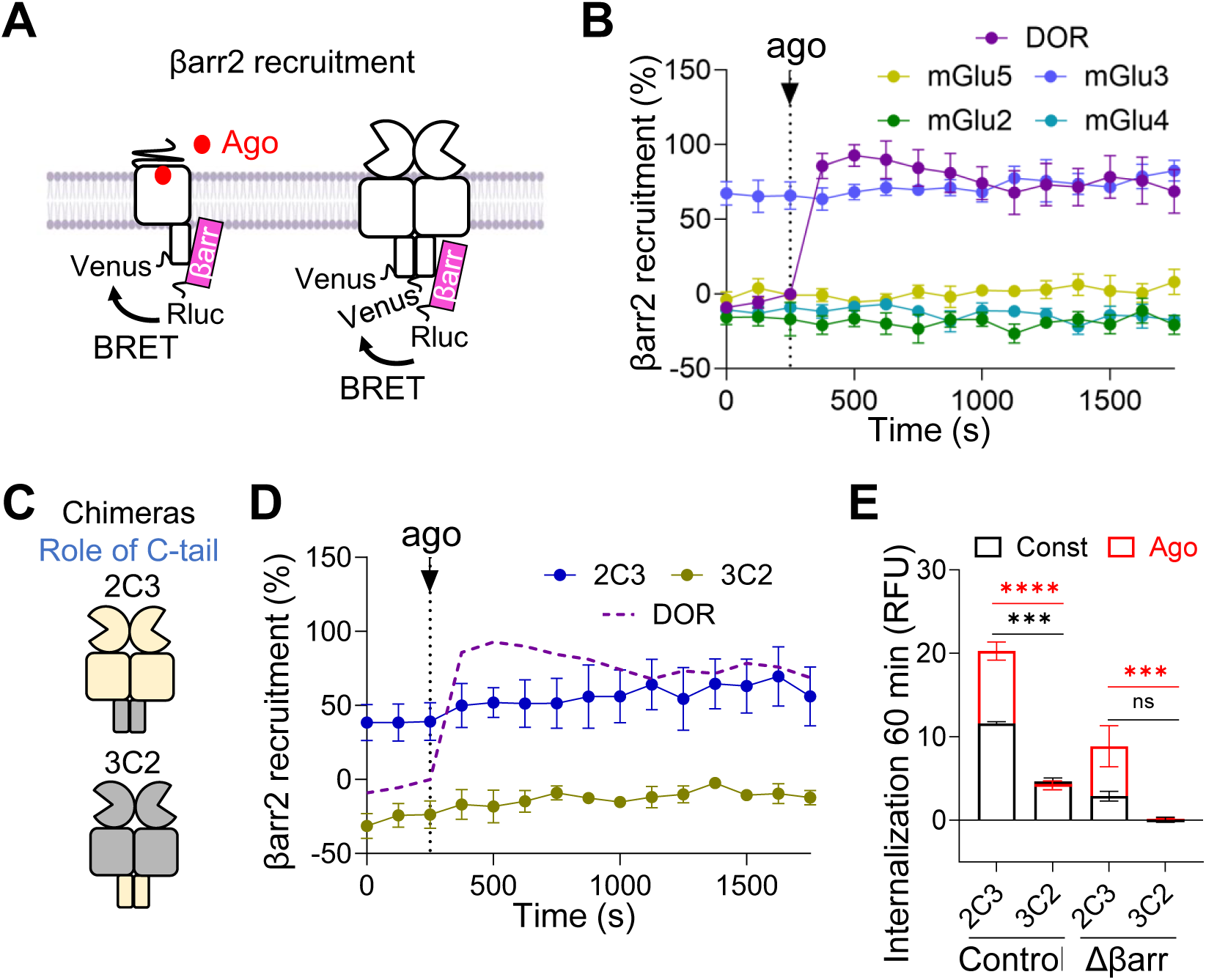
C-terminal domain of mGlu3 is involved in the constitutive association with βarr. (**A**) Schema representing canonical recruitment of βarr and the BRET strategy proposed to evaluate βarr recruitment to GPCRs. (**B**) mGlu-Venus recruitment of βarr2-RLuc measured by BRET before and after addition of a selective agonist (10 μM quisqualate for mGlu5, 10 μM LY354740 for mGlu2 and mGlu3, 10 μM LAP4 for mGlu4). Data are normalized by the maximal mBRET response obtained by the recruitment βarr2-RLuc by DOR-Venus in response to 10 μM snc-162 (100 %) and mBRET values when adding the agonist of DOR (0 %). (**C**) Schema representing 2C3 and 3C2 chimeras’ constructions. (**D**) 2C3-Venus and 3C2-Venus recruitment of βarr2-RLuc measured by BRET before and after addition of a selective agonist (10 μM LY354740). Data are normalized by the maximal response obtained by the recruitment βarr2-RLuc by DOR-Venus in response to 10 μM snc-162. (**E**) Internalization of 2C3 and 3C2 measured after 60 min in the presence of 100 μM LY341495 (constitutive) or agonist-induced by 10 μM LY354740 (+ Ago) in HEK293 cells (control) and Δβarr cells. Data represent the mean ± S.E.M. of at least three independent experiments performed in triplicate.

As GPCRs interact with βarr through the C-terminal domain, we investigated its role in mGlu receptor-βarrestin interaction. We swapped the C-terminal domain of mGlu2 and mGlu3 to build the chimeras 2C3 (mGlu2 with the C-terminal domain of mGlu3) and 3C2 (mGlu3 with the C-terminal domain of mGlu2; Fig. 7C and Fig. S11A). The C-terminal domain of mGlu3 is driving βarrs recruitment as 2C3 keeps basal interaction measured by BRET (Fig. 7D and Fig. S11B). In agreement, 2C3 keeps the internalization properties of mGlu3 either in control cells and Δβarr cells (Fig. 7E and Fig. S12). Also, 3C2 internalizes with the profile of mGlu2 (Fig. 7E and Fig. S13). Altogether, βarrs are required for constitutive internalization of mGlu receptors despite the absence of detectable interaction, except for mGlu3 that constitutively interacts with βarr1 and 2.

### AP2 drives agonist-induced internalization

Finally, we evaluated the role of the tetrameric AP2 protein (α, β, γ, and μ subunits) in mGlu receptor internalization, as AP2 is the final canonical adaptor that binds to βarr and drives endocytosis of agonist-activated GPCRs (Fig. 8A). We used siRNA against the µ AP2 subunit that down regulates AP2 expression (Fig. S14). As previously described, this down regulation largely decreased glucagon-like peptide-1 (GLP1) receptor agonist-induced internalization (Fig. S15)^18^. mGlu3 agonist-induced internalization was almost blunted when down regulating AP2 (Fig. 8B). However, we did not observe any significant effect of AP2 downregulation on the agonist-induced or constitutive internalization of mGlu5, mGlu2 or mGlu4 receptor, neither on the constitutive internalization of mGlu3 (Fig. 8 and Fig. S16).

**Figure 8.**
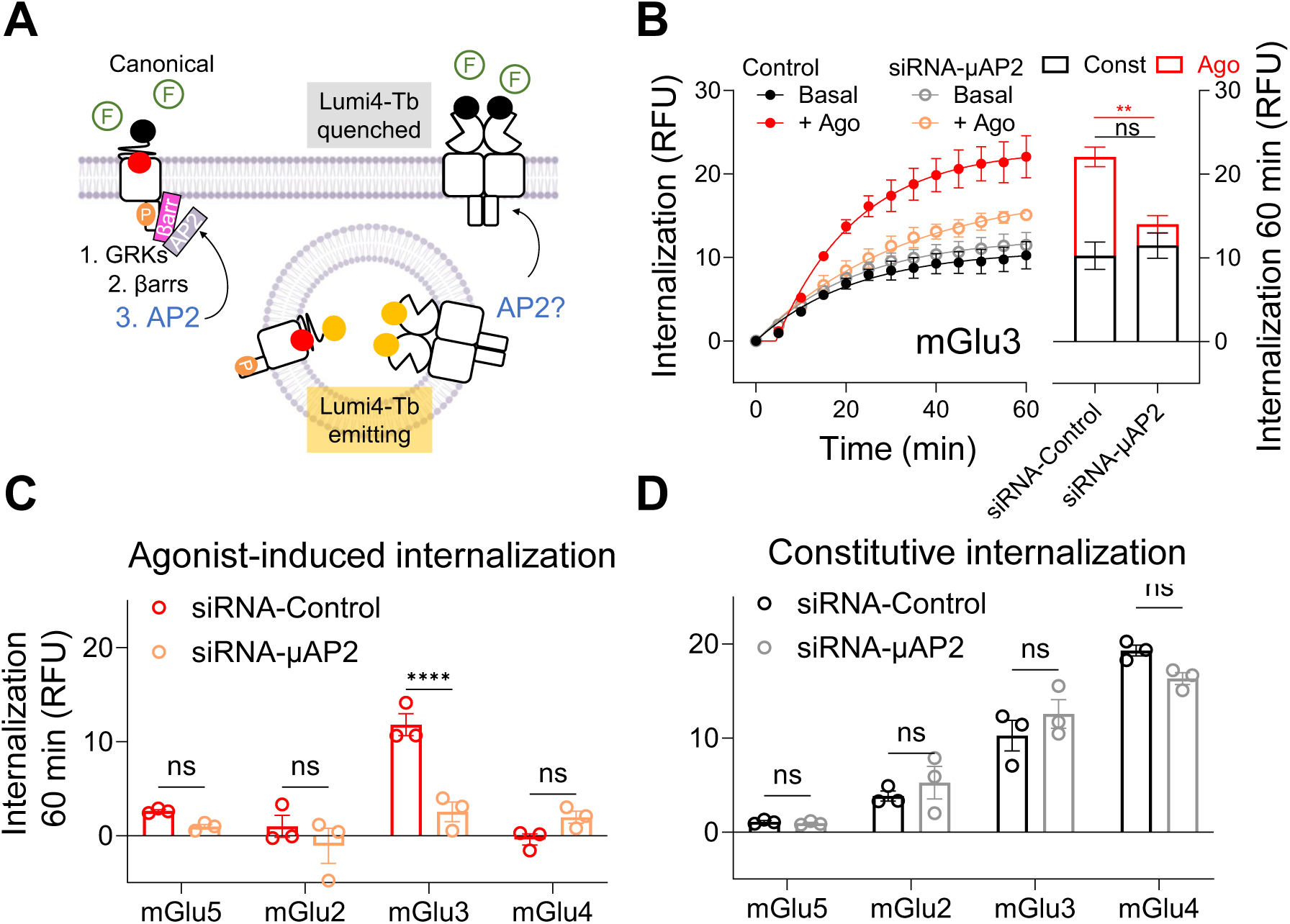
AP2 is required for agonist-induced internalization of mGlu receptors. (**A**) Schema representing canonical impact of AP2 in internalization. (**B**) Internalization of mGlu3 receptor in the absence of agonist (control, 100 μM LY341495) or in the presence of an agonist (+ Ago, 10 μM LY354740), in HEK293 cells (control) versus HEK293 cells treated with siRNA-AP2. Bars represent constitutive internalization (const, given by the basal signal) and agonist induced (ago, calculated by subtracting basal signal) after 60 min of internalization at 37 °C. (**C**) Agonist-induced internalization of mGlu5, mGlu2, mGlu3 and mGlu4 after 60 min of treatment with a selective agonist (10 μM quisqualate for mGlu5, 10 μM LY354740 for mGlu2 and mGlu3, 10 μM LAP4 for mGlu4), in HEK293 cells (red) and in HEK293 cells treated with siRNA-AP2 (orange). (**D**) Constitutive internalization of mGlu5, mGlu2, mGlu3 and mGlu4 after 60 min of treatment with 100 μM LY341495, in HEK293 cells (black), in HEK293 cells treated with siRNA-AP2 (grey). Noteworthy, internalization RFU were lower than in previous experiments as DERET was measured 72 hours after transfection to find the best compromise of siRNA effect and receptor expression. Data represent the mean ± S.E.M. of at least three independent experiments performed in triplicate.

## Discussion

Regulation of GPCR cell surface density is essential for precise physiological function. Highly studied for rhodopsin-like GPCRs, it revealed in most cases the involvement of GRKs, βarrs and AP2 to drive the activated receptor into clathrin coated pits, but other mechanisms are also possible. Surprisingly, not much is known for class C receptors internalization. Our data show that all mGlu receptors are constitutively internalized and only mGlu1, mGlu5 (then likely mGlu1-5), mGlu3 and mGlu2-3, are agonist-induced internalized (Fig. 9). For these receptors, we showed that the agonist-induced internalization follows a non-canonical mechanism, being GRK and βarrs independent, while their constitutive internalization is strictly βarr dependent (Fig. 9).

**Figure 9.**
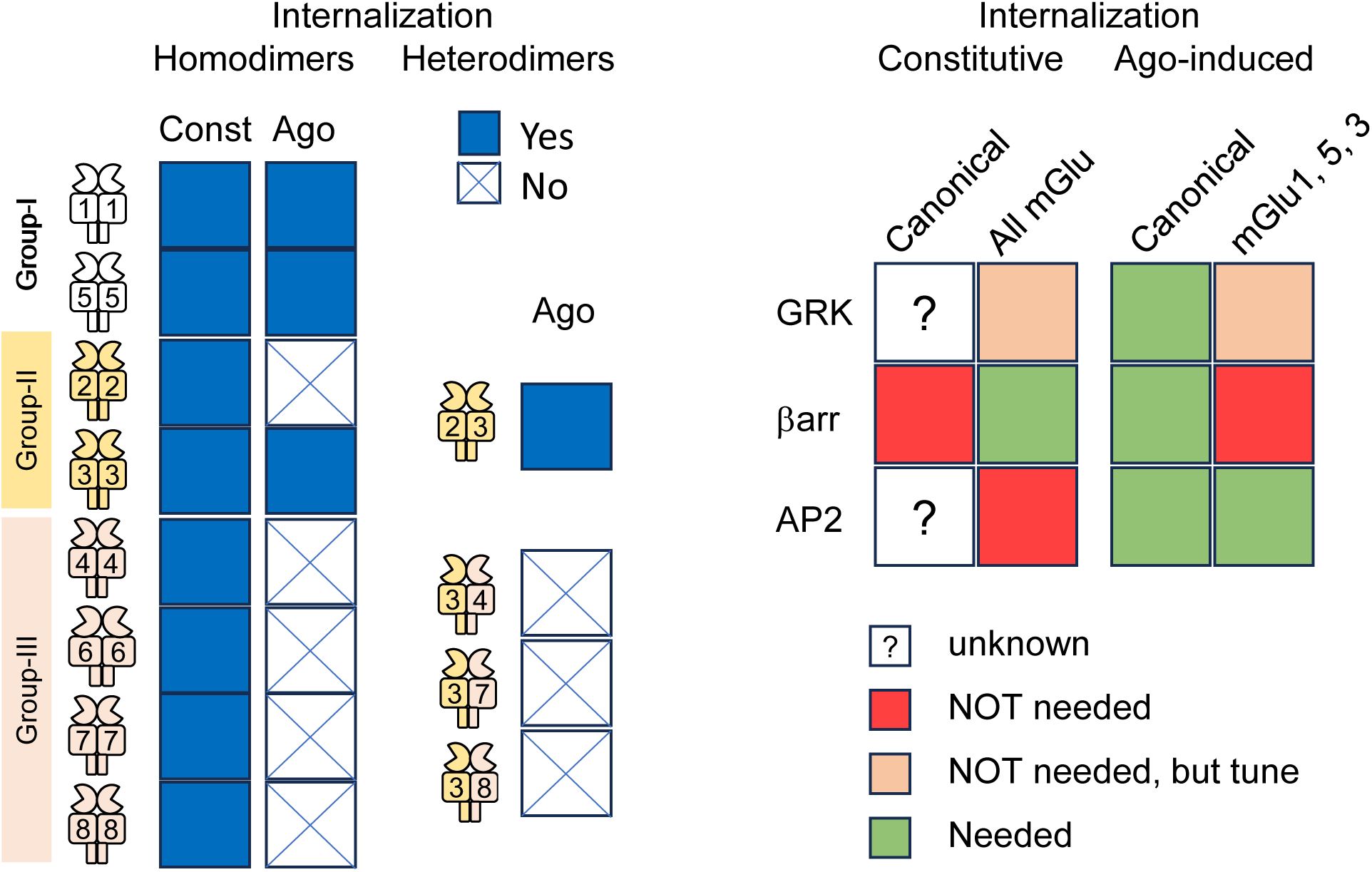
Visual summary of the internalization properties of every mGlu receptors.

Despite years of research on mGlu receptors, how they internalize remains unclear. Yet, some mGlu receptors have been already identified for being internalized in neurons: mGlu1 in Purkinje cells^32^, mGlu5 in enteric^33^ and hippocampal neurons^12^, mGlu7 and mGlu3 also in hippocampal neurons^34,35^. These observations highlight internalization as a process that physiologically regulates mGlu receptor density in the plasma membrane and, thus, their response to drugs.

However, the detection techniques commonly used make the precise determination of mGlu receptor internalization difficult. These include the use of antibodies in living cells, as those can trigger internalization, as demonstrated for MAB1/28 for mGlu7^36^, and thus bias antibody-based internalization measurements. Also, internalization is commonly analysed as an endpoint measurement, where phenomena such as recycling could hide internalization outputs. Indeed, simply measuring membrane receptors may prevent the detection of an internalization process if compensated by insertion of new receptors in the plasma membrane, or receptor recycling from endosomes^9,11^. Moreover, internalized receptors may just stay under the plasma membrane, case described for example for dopamine 1 receptor^37^, and then not resolved as being internalized by microscopy.

To overcome these caveats, measurement of internalization by DERET has been optimized. DERET has been demonstrated to allow the identification of the internalization of several GPCRs, including mGlu5^17^, for which internalization has been described to be allosterically modulated^38,39^. By the use of DERET, we identified both constitutive and agonist-induced internalization of mGlu receptors over time, and ascertain that internalization kinetics are different among mGlu receptors. Specifically, we identified mGlu5 as a receptor that is quickly recycled to the plasma membrane after agonist-induced internalization. Then, time could justify differences observed among receptor internalization outputs in endpoint measurements.

Also, different subtype combinations could drive different profiles of internalization. When evaluating mGlu receptor internalization we showed that only mGlu1, mGlu5 and mGlu3 homodimers are able to be agonist-induced internalized in HEK293 cells, then neither mGlu2 nor any group-III receptors. As such, one is expecting that mGlu1-5 would internalize but not group-III heterodimers upon agonist stimulation. We showed that the mGlu3 subunit can also drive mGlu2 internalization when assembled into a mGlu2-3 heterodimer in intracellular compartments upon agonist stimulation. However, it cannot when associated with mGlu4, mGlu7 or mGlu8. This suggests that the mGlu2 C-terminal domain would not be able to restrict the receptor at the cell surface, in contrast to C-terminal domains of the group-III mGlu receptors. This observation highlights the functional significance of mGlu receptor heterodimer formation^40^ and points out the relevance of considering protein partners when evaluating mGlu receptor internalization.

The use of different cellular models through literature puzzles the accurate interpretation of mGlu receptor internalization. HEK293 cells have a high expression of GRK2 and GRK6^29^, as well as neurons^30^, but GRK4 has been described to participate in endogenous mGlu1 internalization^10^. Also, HEK293 cells lack internalization adaptors that are described for mGlu receptor internalization in neurons, such as PICK-1^35,41^ or norbin^42^. Protein expression could support why we could not detect mGlu7 agonist-induced internalization while it has been described to internalize in neurons^43^.

It was then important to compare the internalization properties of all mGlu receptors in the same cellular background, and using a technique allowing the analysis in real time. Under such condition, we showed that GRKs are dispensable for constitutive and agonist-induced internalization of mGlu receptors. Although not required, they still play a role in the internalization process: only GRK2 and GRK3 can modulate the internalization profile of mGlu5; mGlu2 is specifically agonist-induced internalized likely when phosphorylated by GRK6 but not in the presence of GRK2 and mGlu3 internalization dynamics is affected by GRK overexpression. These GRK-fingerprints for mGlu receptor internalization are not surprising as GRKs are selectively located in the cell: GRK2/3 are soluble in the cytoplasm whereas GRK5/6 are attached to the plasma membrane^44^. GRK availability could determine the efficacity of GRKs in phosphorylating a mGlu receptor to induce internalization, as it happens for opioid receptors^45^. The role of GRK4 has not been studied as HEK293 cells are not expressing this GRK subtype^29^. Future work will be needed to ascertain the role of GRK4 in mGlu receptor internalization, as its role in mGlu desensitization has been reported^10^.

In contrast to canonical GPCR internalization mechanism, βarrs are not needed for the agonist-induced internalization of mGlu1, 5 and 3, and no βarrs recruitment could be observed on these receptors upon activation, in agreement with previous reports^46,47^. Although βarrs have been reported to be involved in the mGlu1 internalization using a dominant negative construct, this study did not separate the constitutive internalization that we showed is βarr dependent, and agonist-induced internalization^10^. Of interest, βarrs are known to bind within the cavity created by the movement of transmembrane domain 6 (TM6) in activated class A GPCRs^48^, a cavity that cannot exist in class C GPCRs as no movement of TM6 is observed upon activation^49,50^.

However, βarrs are required for constitutive internalization of mGlu receptors. Same as for another class C GPCR, the GABA_B_ receptor, which has been described to be constitutively internalized in a clathrin-dependent manner in neurons^51^. The mGlu3 receptor is unique, as it is the only one constitutively interacting with βarrs, as V1b^52^, ACKR4^53^ or CCR1^54^. This could explain the role of βarrs in the constitutive internalization of mGlu3. Yet, no constitutive close proximity with βarrs could be detected for the other mGlu receptors, even though they also show strong constitutive internalization, suggesting an indirect role of βarrs for the constitutive internalization of mGlu receptors that needs to be investigated.

Due to the fact that βarrs are not participating in mGlu receptor agonist-induced internalization, AP2 should not play a role according to the canonical mechanisms of internalization. Intriguingly, AP2 did not play a role in constitutive internalization of mGlu receptors but it does for agonist-induced internalization of mGlu3. mGlu3 receptors emerge as possible direct recruiter of AP2 as observed with GLP1^18^, an interaction that may not need βarrs.

In this work, we precisely identified the constitutive and the agonist-induced mechanisms of mGlu receptors in HEK293 cells. In this cellular background, constitutive internalization relies in βarrs whereas mGlu3 agonist-induced internalization is dependent on AP2. GRKs appear as modulators of the kinetic profiles of these mGlu receptors. Nevertheless, mGlu receptors could follow diverse internalization mechanisms in neurons as it has been already suggested elsewhere for mGlu5^55^. This study provides a solid foundation for the evaluation of internalization of mGlu receptors in different cell types, such as different brain cells.

## Materials and methods

### Reagents

LY341495 disodium salt (4062), L-Quisqualic acid (0188), LY354740 (3246), DAMGO (1171), exendin-4 (1933) and L-AP4 (0130) were purchased from Tocris Bioscience (Noyal Châtillon sur Seiche, France). Lumi-4-Tb labelling reagents, lumi-4-Tb labelled antibodies and Tag-lite SNAP/CLIP Labeling Medium (#LABMED, Revvity) were kindly supplied by Revvity. DMEM (41965-039), DMEM + GlutaMAX™-I (31966-021) and Opti-MEM® (31985-047) were purchased from Gibco (Thermo Fisher Scientific, Darmstadt, Germany). Lipofectamine 2000 (11668019) was obtained from Life Technologies (Carlsbad, CA, USA). Coelenterazine-h (S2011) was purchased from Promega Corporation (Madison, WI, USA). SNAP-Surface® Block (S9143S) was purchased from New England Biolabs (Ipswich, MA, USA). All other reagents were purchased in Sigma Aldrich (Munich, Germany).

### Plasmids

Plasmids encoding GPCRs were FLAG-Halo, FLAG-Clip or FLAG-SNAP tagged in the N-terminal domain of the receptor and already available in the laboratory. Venus-fluorescent protein was added in the C-terminal domain for DOR, mGlu5, mGlu2, mGlu3, mGlu4, 2C3 and 3C2 by Gene Cust (Boynes, France). β-arrestins were RLuc8 tagged in the C-terminal domain. Otherwise stated, constructs were human and inserted in pcDNA3.1 backbones.

### Cell lines

Unless specified, HEK 293 cells (CRL-1573™, ATCC) were used. Δβarr1/2 HEK 293 cell line was a gift from Dr. Inoue (Tohoku University, Japan); ΔQ-GRK HEK 293 cell line was a gift from Dr. Hoffman (Jena University, Germany). Cells were maintained in DMEM supplemented with 10 % FBS and 2 % penicillin/streptomycin (ΔβArr1/2 and ΔQ-GRK cells) at 37 °C and 5 % CO_2_, and used up to passage 30. All cell lines were tested negative for *Mycoplasma* contamination.

Cells were transiently transfected in Opti-MEM (Gibco, Thermo Fisher Scientific) using lipofectamine 2000 (Thermo Fisher Scientific). When transfecting mGlu receptors, the glutamate transporter EAAC1 was co-transfected to reduce the amount of ambient glutamate in the extracellular medium. The total amount of transfected DNA was kept equal by supplementing the transfection mix with the empty plasmid.

### Silencing of protein expression

Cells were plated at a density of 50000 cells/well and reverse-transfected with DharmaFECT1 Transfection reagent (Horizon), the plasmid encoding the GPCR under study and 50 nM of control-siRNA or 50 nM ON-TARGETplus Human AP2-µ2subunit (Horizon). Experiments were performed 72 h after.

### DERET (internalization assay)

Cells were seeded at a density between 50000 and 100000 cells/well depending on the cell type and reverse-transfected with lipofectamine and the targeted receptors labelled with a suicide enzyme, SNAP- or Halo-tag. 24 h after and prior to the experiment, cell medium was changed with medium containing GlutaMAX for 90 min at 37 °C, for reducing the amount of ambient glutamate during the experiment.

Then, cells were treated with 100 nM of Tag-lite SNAP-Lumi4-Tb Labeling Reagent (#SSNPTBD, Revvity) or 100 nM of Tag-lite HaloTag-Lumi4-Tb Labeling Reagent (#SHALOTBC, Revvity) labelling reagent for 1 h at 4 °C in 1x Tag-lite SNAP/CLIP Labeling Medium (#LABMED, Revvity). Cells were washed with 100 µl of 1x Tag-lite SNAP/CLIP Labeling Medium (#LABMED, Revvity) twice and receptor expression was measured in Spark® multimode microplate reader (Tecan). Compounds were then added at the indicated concentrations together with 25 µM fluorescein. The signal at 535 nm and 620 nm was recorded every 5 min during 60 min at 37 °C in Spark® multimode microplate reader (Tecan). In case of pharmacologically inhibiting the phosphorylation of GRK2/3 with cmpd101, after labelling with Tag-lite SNAP-Lumi4-Tb Labeling Reagent (#SSNPTBD, Revvity), cells were treated with 10 μM cmpd101 for 30 min at 4 °C.

Kinetic data was expressed as internalization units in random fluorescent units (RFU), following the equation:

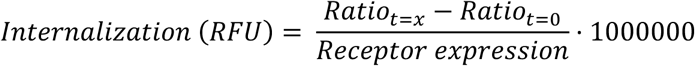

where ratio refers to the emission at 620 nm (integrated during 1500 μs with a delay of measure of 1500 μs) over the emission at 535 nm (integrated during 150 μs with a delay of measure of 50 μs). Receptor expression refers to the emission signal at 620 nm before addition of fluorescein.

Data in bars was expressed as internalization units in RFU following the same formula as stated in kinetics data. Constitutive internalization was the internalization signal obtained after treating cells for 60 min with vehicle; agonist-induced internalization resulted from the signal obtained after agonist-treatment during 60 min minus the signal obtained for constitutive internalization. Otherwise noted, the vehicle condition is 100 μM L341495, to avoid the effect of ambient glutamate.

### Recycling measurements (DERET)

Recycling measurement were DERET-based and performed 60 min after agonist-induced internalization at 37 °C. Then, the agonist was removed (washed) and internalization measurements were done after 120 min, 180 min, 240 min and 300 min in the presence of 25 µM fluorescein. For antagonist accelerated recycling, after agonist washing 10 μM LY341495 were added. Measurements were done after 120 min, 180 min, 240 min and 300 min in the presence of 25 µM fluorescein. Ago-internalization represent the specific internalization signal by subtraction of constitutive internalization. All recordings were done in Spark® multimode microplate reader (Tecan).

### TR-FRET for heterodimer detection

Cells were plated at a density of 100000 cells/well and reverse transfected with two different mGlu receptors, each tagged with a different suicide: either SNAP, CLIP or Halo. 24 h after, cells were incubated either with a Tag-lite SNAP-Lumi4-Tb Labeling Reagent (#SSNPTBD, Revvity), Tag-lite CLIP-Green Labeling Reagent (#SCLPGRNE, Revvity) or Tag-lite HaloTag-Green Labeling Reagent (#SHALOGRNE, Revvity)substrate or a combination of both at a concentration of 100 nM for 1 h at 37 °C in 1X Tag-lite SNAP/CLIP Labeling Medium (#LABMED, Revvity). Then, cells were washed 3 times with 100 μl 1X Tag-lite SNAP/CLIP Labeling Medium (#LABMED, Revvity) and TR-FRET was measured in PheraStar microplate reader (BMG LabTech).

### BRET for βarr recruitment detection

Cells were plated at density of 100000 cells/well and reverse transfected: with the indicated GPCR tagged with a Venus protein for measuring direct recruitment. The indicated βarr tagged with RLuc8 was co-expressed in a proportion 3 to 1 (GPCR:βarr). After 24 h, the medium was changed with medium containing GlutaMAX for 90 min at 37 °C. Cells were washed with PBS twice and incubated 10 min at 37 °C. Receptor expression was measured in Mithras LB 940 Multimode Microplate Reader (Berthold Technologies) by measuring fluorescence emitted at 535 nm. Then, 5 µM coelenterazine was added and BRET measurements were done during 500 s, before the indicated compounds were added. BRET signal was then measured for 1500 s. miliBRET was calculated as the signal obtained at 535 nm over the signal obtained at 485 nm and adjusted by subtracting the ratios obtained when RLuc fusion proteins were expressed alone, and by multiplying the result by 1000. Data was corrected by the BRET signal obtained as a result of DOR recruitment of βarr, and represented then as % of βarr2 recruitment. Bars represent the % of the peak obtained just after addition of an agonist. Otherwise noted, the vehicle condition is 100 μM L341495, to avoid the effect of ambient glutamate.

### Western blot

Cells plated in 100 mm Petri dishes were washed once with ice-cold PBS and subsequently lysed with RIPA Buffer (ThermoFisher, 89900), supplemented with protease inhibitor cocktails (Merck, 5892953001). Cleared lysates were boiled with sodium dodecyl sulphate (SDS) loading buffer and 15 μg of total protein were loaded onto each lane of 8 % polyacrylamide gels. After transfer onto nitrocellulose membranes, the total protein was detected by using specific antibodies (AP2: mouse anti-AP2 m2, BD Biosciences, 611351, (1:250), Actin: mouse anti-b actin, Sigma-Aldrich, A5441(1:5000)). As secondary antibodies, rabbit HTRF pAb Anti-Mouse IgG d2-Conjugate (#61PAMDAA, Revvity) (1:100) was used.

### Statistical analysis

All data analysis was performed using Prism 10 (GraphPad Software Inc., San Diego, CA, USA). Experiments were performed in triplicate, and final experiments represent the mean of at least three independent experiments. Statistical significance was assessed by an ordinary two-way ANOVA followed by: Šídák’s multiple comparisons test, in case of comparison of two different groups; or by Tukey’s multiple comparisons test, in case of comparison of different conditions in the same group. Internalization kinetic data was adjusted to the pharmacokinetic equations described by Hoare S et al., 2020.

### Resource availability

Datasets and metadata will be available in the repository Zenodo. Other information or materials will be available under request.

## Acknowledgements

We thank all the members of Rondard’s Team for the fruitful discussions. We thank the Arpège platform of the Institute of Functional Genomics for providing facilities and technical support and Revvity for providing reagents. M.C. was supported by a postdoctoral fellowship granted by the Alfonso Martin Escudero Foundation (FAME) and a Marie Sklodowska-Curie Individual Fellowship from the European Union’s Horizon 2020 research and innovation programme to the project GLUGLU (project number 101065171). M.H. was recipient of a Doctoral fellowship from the University of Montpellier (2024). P.-A.L. was the recipient of a postdoctoral fellowship from the LabEX MAbImprove (grant NR-10-LABX-5301). This work was supported by Fondation pour la Recherche Médicale (DEQ20170336747 to JPP, EQU202303016470 to PR), the Agence Nationale pour la Recherche (ANR-22-CE16-0002-01 to JPP) and Revvity (Codolet, France).

## Author contributions

M.C. conceptualized the project, performed experiments, data analysis and wrote the manuscript. J.L. performed the Western blot. D.M. and I.B. set the basis with preliminary data. J.D., C.H. and A.I. provided the cell lines. M.H. provided experimental support. P.R. got funding for the project. P-A. L gave critical review and constructive discussions. J-P.P and L.P. conceptualized the project, supervised the work and wrote the manuscript. All authors approved the manuscript.

## Supplemental information

Document S1: Figures S1-S16 and Table S1

## References

1. Hauser AS, Attwood MM, Rask-Andersen M, Schiöth HB, Gloriam DE. Trends in GPCR drug discovery: new agents, targets and indications. Nat Rev Drug Discov 2017 1612. 2017;16(12):829–842. doi:10.1038/nrd.2017.178

2. Kaksonen M, Roux A. Mechanisms of clathrin-mediated endocytosis. Nat Rev Mol Cell Biol 2018 195. 2018;19(5):313–326. doi:10.1038/nrm.2017.132

3. Moo E Von, Senten JR van, Bräuner-Osborne H, Møller TC. Arrestin-Dependent and - Independent Internalization of G Protein–Coupled Receptors: Methods, Mechanisms, and Implications on Cell Signaling. Mol Pharmacol. 2021;99(4):242–255. doi:10.1124/MOLPHARM.120.000192

4. Tian X, Kang DS, Benovic JL. β-arrestins and G Protein-Coupled Receptor Trafficking. Handb Exp Pharmacol. 2014;219:173. doi:10.1007/978-3-642-41199-1_9

5. Scarselli M, Donaldson JG. Constitutive internalization of G protein-coupled receptors and G proteins via clathrin-independent endocytosis. J Biol Chem. 2009;284(6):3577–3585. doi:10.1074/JBC.M806819200

6. Spiess K, Bagger SO, Torz LJ, et al. Arrestin-independent constitutive endocytosis of GPR125/ADGRA3. Ann N Y Acad Sci. 2019;1456(1):186–199. doi:10.1111/NYAS.14263

7. Iacovelli L, Nicoletti F, De Blasi A. Molecular mechanisms that desensitize metabotropic glutamate receptor signaling: an overview. Neuropharmacology. 2013;66:24–30. doi:10.1016/J.NEUROPHARM.2012.05.005

8. Ribeiro FM, Ferreira LT, Paquet M, et al. Phosphorylation-independent regulation of metabotropic glutamate receptor 5 desensitization and internalization by G protein-coupled receptor kinase 2 in neurons. J Biol Chem. 2009;284(35):23444–23453. doi:10.1074/JBC.M109.000778

9. Abreu N, Acosta-Ruiz A, Xiang G, Levitz J. Mechanisms of differential desensitization of metabotropic glutamate receptors. Cell Rep. 2021;35(4). doi:10.1016/J.CELREP.2021.109050

10. Mundell SJ, Pula G, McIlhinney RAJ, Roberts PJ, Kelly E. Desensitization and internalization of metabotropic glutamate receptor 1a following activation of heterologous Gq/11-coupled receptors. Biochemistry. 2004;43(23):7541–7551. doi:10.1021/BI0359022

11. Lee J, Gonzalez-Hernandez A, Kristt M, et al. Distinct beta-arrestin coupling and intracellular trafficking of metabotropic glutamate receptor homo- and heterodimers. Sci Adv. 2023;9(49). doi:10.1126/SCIADV.ADI8076

12. Fourgeaud L, Bessis A-S, oise Rossignol F, Pin J-P, Olivo-Marin J-C, Hé mar A. The Metabotropic Glutamate Receptor mGluR5 Is Endocytosed by a Clathrin-independent Pathway*. Published online 2003. doi:10.1074/jbc.M205663200

13. Møller TC, Moo E Von, Inoue A, Pedersen MF, Bräuner-Osborne H. Characterization of the real-time internalization of nine GPCRs reveals distinct dependence on arrestins and G proteins. Biochim Biophys Acta - Mol Cell Res. 2024;1871(1):119584. doi:10.1016/J.BBAMCR.2023.119584

14. Kaur Dhami G, Anborgh PH, Dale LB, Sterne-Marr R, Ferguson SSG. Phosphorylation-independent Regulation of Metabotropic Glutamate Receptor Signaling by G Protein-coupled Receptor Kinase 2*. J Biol Chem. 2002;277:25266–25272. doi:10.1074/jbc.M203593200

15. Crupi R, Impellizzeri D, Cuzzocrea S. Role of Metabotropic Glutamate Receptors in Neurological Disorders. Front Mol Neurosci. 2019;12:20. doi:10.3389/FNMOL.2019.00020

16. Adams DH, Zhang L, Millen BA, Kinon BJ, Gomez J-C. Pomaglumetad Methionil (LY2140023 Monohydrate) and Aripiprazole in Patients with Schizophrenia: A Phase 3, Multicenter, Double-Blind Comparison. Schizophr Res Treatment. 2014;2014:758212. doi:10.1155/2014/758212

17. Levoye A, Zwier JM, Jaracz-Ros A, et al. A broad G protein-coupled receptor internalization assay that combines SNAP-tag labeling, diffusion-enhanced resonance energy transfer, and a highly emissive terbium cryptate. Front Endocrinol (Lausanne). 2015;6(NOV):167. doi:10.3389/fendo.2015.00167

18. Liu J, Xue L, Ravier M, et al. Multi-faceted roles of β-arrestins in G protein-coupled receptors endocytosis. bioRxiv. Published online January 20, 2024:2024.01.18.576020. doi:10.1101/2024.01.18.576020

19. Hoare SRJ, Tewson PH, Quinn AM, Hughes TE, Bridge LJ. Analyzing kinetic signaling data for G-protein-coupled receptors. Sci Reports 2020 101. 2020;10(1):1–23. doi:10.1038/s41598-020-67844-3

20. Acher F. Metabotropic glutamate receptors. Tocris Biosci Rev Lett. 2011;26:1–10. Accessed January 16, 2025. https://resources.tocris.com/pdfs/literature/reviews/mglur-review-2019-web.pdf

21. Tora AS, Rovira X, Cao AM, et al. Chloride ions stabilize the glutamate-induced active state of the metabotropic glutamate receptor 3. Neuropharmacology. 2018;140:275–286. doi:10.1016/j.neuropharm.2018.08.011

22. Doumazane E, Scholler P, Zwier JM, Trinquet E, Rondard P, Pin J. A new approach to analyze cell surface protein complexes reveals specific heterodimeric metabotropic glutamate receptors. FASEB J. 2011;25(1):66–77. doi:10.1096/fj.10-163147

23. Kniazeff J, Bessis AS, Maurel D, Ansanay H, Prézeau L, Pin JP. Closed state of both binding domains of homodimeric mGlu receptors is required for full activity. Nat Struct Mol Biol 2004 118. 2004;11(8):706–713. doi:10.1038/nsmb794

24. von Zastrow M, Williams JT. Modulating neuromodulation by receptor membrane traffic in the endocytic pathway. Neuron. 2012;76(1):22. doi:10.1016/J.NEURON.2012.09.022

25. Perkovska S, Méjean C, Ayoub MA, et al. V1b vasopressin receptor trafficking and signaling: Role of arrestins, G proteins and Src kinase. Traffic. 2018;19(1):58–82. doi:10.1111/tra.12535

26. Meng J, Xu C, Lafon PA, et al. Nanobody-based sensors reveal a high proportion of mGlu heterodimers in the brain. Nat Chem Biol. 2022;18(8):894–903. doi:10.1038/S41589-022-01050-2

27. TC M, MF P, JR van S, et al. Dissecting the roles of GRK2 and GRK3 in μ-opioid receptor internalization and β-arrestin2 recruitment using CRISPR/Cas9-edited HEK293 cells. Sci Rep. 2020;10(1). doi:10.1038/S41598-020-73674-0

28. Drube J, Haider RS, Matthees ESF, et al. GPCR kinase knockout cells reveal the impact of individual GRKs on arrestin binding and GPCR regulation. Nat Commun. 2022;13(1). doi:10.1038/S41467-022-28152-8

29. Reichel M, Weitzel V, Klement L, Hoffmann C, Drube J. Suitability of GRK Antibodies for Individual Detection and Quantification of GRK Isoforms in Western Blots. Int J Mol Sci. 2022;23(3). doi:10.3390/IJMS23031195

30. Guimarães TR, Swanson E, Kofler J, Thathiah A. G protein-coupled receptor kinases are associated with Alzheimer’s disease pathology. Neuropathol Appl Neurobiol. 2021;47(7):942–957. doi:10.1111/NAN.12742

31. Alvarez-Curto E, Inoue A, Jenkins L, et al. Targeted Elimination of G Proteins and Arrestins Defines Their Specific Contributions to Both Intensity and Duration of G Protein-coupled Receptor Signaling. J Biol Chem. 2016;291(53):27147–27159. doi:10.1074/JBC.M116.754887

32. Sallese M, Salvatore L, D’Urbano E, et al. The G-protein-coupled receptor kinase GRK4 mediates homologous desensitization of metabotropic glutamate receptor 1. FASEB J. 2000;14(15):2569–2580. doi:10.1096/FJ.00-0072COM

33. Liu MT, Kirchgessner AL. Agonist- and Reflex-Evoked Internalization of Metabotropic Glutamate Receptor 5 in Enteric Neurons. J Neurosci. 2000;20(9):3200. doi:10.1523/JNEUROSCI.20-09-03200.2000

34. Pelkey KA, Yuan X, Lavezzari G, Roche KW, McBain CJ. mGluR7 undergoes rapid internalization in response to activation by the allosteric agonist AMN082. Neuropharmacology. 2007;52(1):108–117. doi:10.1016/J.NEUROPHARM.2006.07.020

35. Tuduri P, Bouquier N, Girard B, et al. Modulation of Hippocampal Network Oscillation by PICK1-Dependent Cell Surface Expression of mGlu3 Receptors. J Neurosci. 2022;42(47):8897–8911. doi:10.1523/JNEUROSCI.0063-22.2022

36. Ullmer C, Zoffmann S, Bohrmann B, et al. Functional monoclonal antibody acts as a biased agonist by inducing internalization of metabotropic glutamate receptor 7. Br J Pharmacol. 2012;167(7):1448–1466. doi:10.1111/j.1476-5381.2012.02090.x

37. Kotowski SJ, Hopf FW, Seif T, Bonci A, von Zastrow M. Endocytosis Promotes Rapid Dopaminergic Signaling. Neuron. 2011;71(2):278–290. doi:10.1016/J.NEURON.2011.05.036/ATTACHMENT/D038C525-3485-4245-9904-BC0414BE444F/MMC4.MOV

38. Arsova A, Møller TC, Vedel L, et al. Detailed in Vitro Pharmacological Characterization of Clinically Tested Negative Allosteric Modulators of the Metabotropic Glutamate Receptor 5. Mol Pharmacol. 2020;98(1):49–60. doi:10.1124/mol.119.119032

39. Arsova A, Møller TC, Hellyer SD, et al. Positive allosteric modulators of metabotropic glutamate receptor 5 as tool compounds to study signaling bias. Mol Pharmacol. Published online February 18, 2021:MOLPHARM-AR-2020-000185. doi:10.1124/molpharm.120.000185

40. Belkacemi K, Rondard P, Pin JP, Prézeau L. Heterodimers Revolutionize the Field of Metabotropic Glutamate Receptors. Neuroscience. Published online 2024. doi:10.1016/J.NEUROSCIENCE.2024.06.013

41. Ramsakha N, Ojha P, Pal S, Routh S, Citri A, Bhattacharyya S. A vital role for PICK1 in the differential regulation of metabotropic glutamate receptor internalization and synaptic AMPA receptor endocytosis. J Biol Chem. 2023;299(6):104837. doi:10.1016/J.JBC.2023.104837

42. Ojha P, Pal S, Bhattacharyya S. Regulation of Metabotropic Glutamate Receptor Internalization and Synaptic AMPA Receptor Endocytosis by the Postsynaptic Protein Norbin. J Neurosci. 2022;42(5):731–748. doi:10.1523/JNEUROSCI.1037-21.2021

43. Lavezzari G, Roche KW. Constitutive endocytosis of the metabotropic glutamate receptor mGluR7 is clathrin-independent. Neuropharmacology. 2007;52(1):100–107. doi:10.1016/J.NEUROPHARM.2006.07.011

44. Ribas C, Penela P, Murga C, et al. The G protein-coupled receptor kinase (GRK) interactome: Role of GRKs in GPCR regulation and signaling. Biochim Biophys Acta - Biomembr. 2007;1768(4):913–922. doi:10.1016/J.BBAMEM.2006.09.019

45. Jullié D, Benitez C, Knight TA, Simic MS, von Zastrow M. Endocytic trafficking determines cellular tolerance of presynaptic opioid signaling. Elife. 2022;11. doi:10.7554/ELIFE.81298

46. Dhami GK, Ferguson SSG. Regulation of metabotropic glutamate receptor signaling, desensitization and endocytosis. Pharmacol Ther. 2006;111(1):260–271. doi:10.1016/J.PHARMTHERA.2005.01.008

47. Pin JP, Bettler B. Organization and functions of mGlu and GABA B receptor complexes. Nature. 2016;540(7631):60–68. doi:10.1038/nature20566

48. Rasmussen SGF, Devree BT, Zou Y, et al. Crystal structure of the β2 adrenergic receptor–Gs protein complex. Nat 2011 4777366. 2011;477(7366):549–555. doi:10.1038/nature10361

49. Shen C, Mao C, Xu C, et al. Structural basis of GABAB receptor–Gi protein coupling. Nature. Published online April 28, 2021:1–5. doi:10.1038/s41586-021-03507-1

50. Lin S, Han S, Cai X, et al. Structures of Gi-bound metabotropic glutamate receptors mGlu2 and mGlu4. Nat 2021 5947864. 2021;594(7864):583–588. doi:10.1038/s41586-021-03495-2

51. Vargas KJ, Terunuma M, Tello JA, Pangalos MN, Moss SJ, Couve A. The Availability of Surface GABA B Receptors Is Independent of-Aminobutyric Acid but Controlled by Glutamate in Central Neurons * □ S. Published online 2008. doi:10.1074/jbc.M802419200

52. Koshimizu T, Honda K, Nagaoka-Uozumi S, et al. Complex formation between the vasopressin 1b receptor, β-arrestin-2, and the μ-opioid receptor underlies morphine tolerance. Nat Neurosci 2018 216. 2018;21(6):820–833. doi:10.1038/s41593-018-0144-y

53. Matti C, Salnikov A, Artinger M, et al. ACKR4 Recruits GRK3 Prior to β-Arrestins but Can Scavenge Chemokines in the Absence of β-Arrestins. Front Immunol. 2020;11. doi:10.3389/fimmu.2020.00720

54. Gilliland CT, Salanga CL, Kawamura T, Trejo J, Handel TM. The chemokine receptor CCR1 is constitutively active, which leads to G protein-independent, β-arrestin-mediated internalization. J Biol Chem. 2013;288(45):32194–32210. doi:10.1074/JBC.M113.503797

55. Scheefhals N, Catsburg LAE, Westerveld ML, Blanpied TA, Hoogenraad CC, MacGillavry HD. Shank Proteins Couple the Endocytic Zone to the Postsynaptic Density to Control Trafficking and Signaling of Metabotropic Glutamate Receptor 5. Cell Rep. 2019;29(2):258–269.e8. doi:10.1016/j.celrep.2019.08.102

