## Supplementary material for "Non-canonical internalization mechanisms of mGlu receptors": supp figures

### Supplemental information


**Figure S1**. Control of mGlu5 receptor expression in HEK293 cells.

**Figure S2.** Control of mGlu5-Y receptor expression in HEK293 cells.

**Figure S3.** Control mGlu receptor expression in HEK293 cells.

**Figure S4.** Internalization kinetics of mGlu receptors.

**Figure S5.** Kinase-dead GRKs impact on mGlu3 receptor internalization.

**Figure S6.** Effect of GRK3 and GRK6 on mGlu receptor internalization.

**Figure S7.** Control of mGlu receptor expression in Δβarr cells.

**Figure S8.** Internalization kinetics of mGlu receptors in cells lacking βarrs (Δβarr) and in Δβarr cells supplemented with βarr2 (+ βarr2).

**Figure S9.** βarr1 effect on mGlu receptor internalization.

**Figure S10.** Control of BRET signal for βarr2 recruitment and effect of βarr1.

**Figure S11.** C-terminal exchange between mGlu2 and mGlu3.

**Figure S12.** 2C3 internalization profiles.

**Figure S13.** 3C2 internalization profile.

**Figure S14.** Detection of Western blot of the μ-subunit of AP2 in HEK293 cells.

**Figure S15.** Internalization kinetics of GLP-1 measured in HEK293 cells (control) and HEK293 cells treated with siRNA-μAP2.

**Figure S16.** Internalization kinetics of mGlu5, mGlu2 and mGlu4 measured in HEK293 cells (control) and HEK293 cells treated with siRNA-μAP2.

**Table S1**. Pharmacological parameters of internalization of mGlu5.

### Fig S1

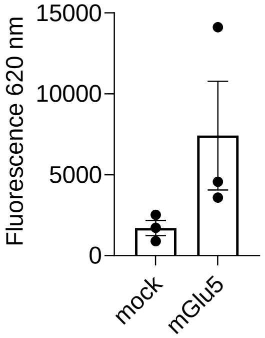


**Figure S1.** Control of mGlu5 receptor expression in HEK293 cells. mGlu5 receptor expression measured by SNAP-lumi4-Tb labelling for 1 h at 4 ºC in comparison with non-transfected cells (mock). Data represent the mean ± S.E.M. of three independent experiments performed in triplicate.

#### Fig S2

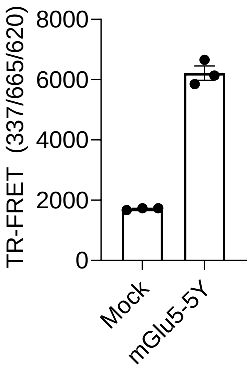


**Figure S2.** Control of mGlu5-Y receptor expression in HEK293 cells. mGlu5-Y receptor expression measured by TR-FRET between CLIP-mGlu5 tagged with SNAP-lumi4-Tb and SNAP-mGlu5Y tagged with CLIP-green in comparison with non-transfected cells (mock). Data represent the mean ± S.E.M. of three independent experiments performed in triplicate.

### Fig S3

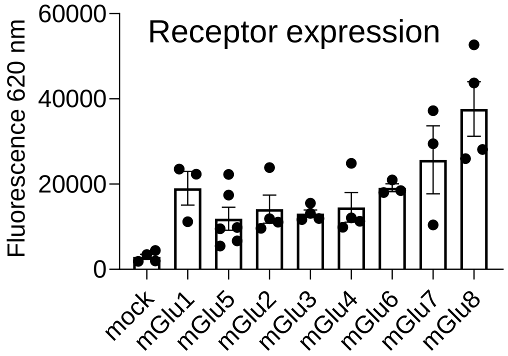


**Figure S3.** Control mGlu receptor expression in HEK293 cells. mGlu receptor expression measured by SNAP-lumi4-Tb labelling for 1 h at 4 ºC in comparison with non-transfected cells (mock). Data represent the mean ± S.E.M. of at least three independent experiments performed in triplicate.

Fig S4

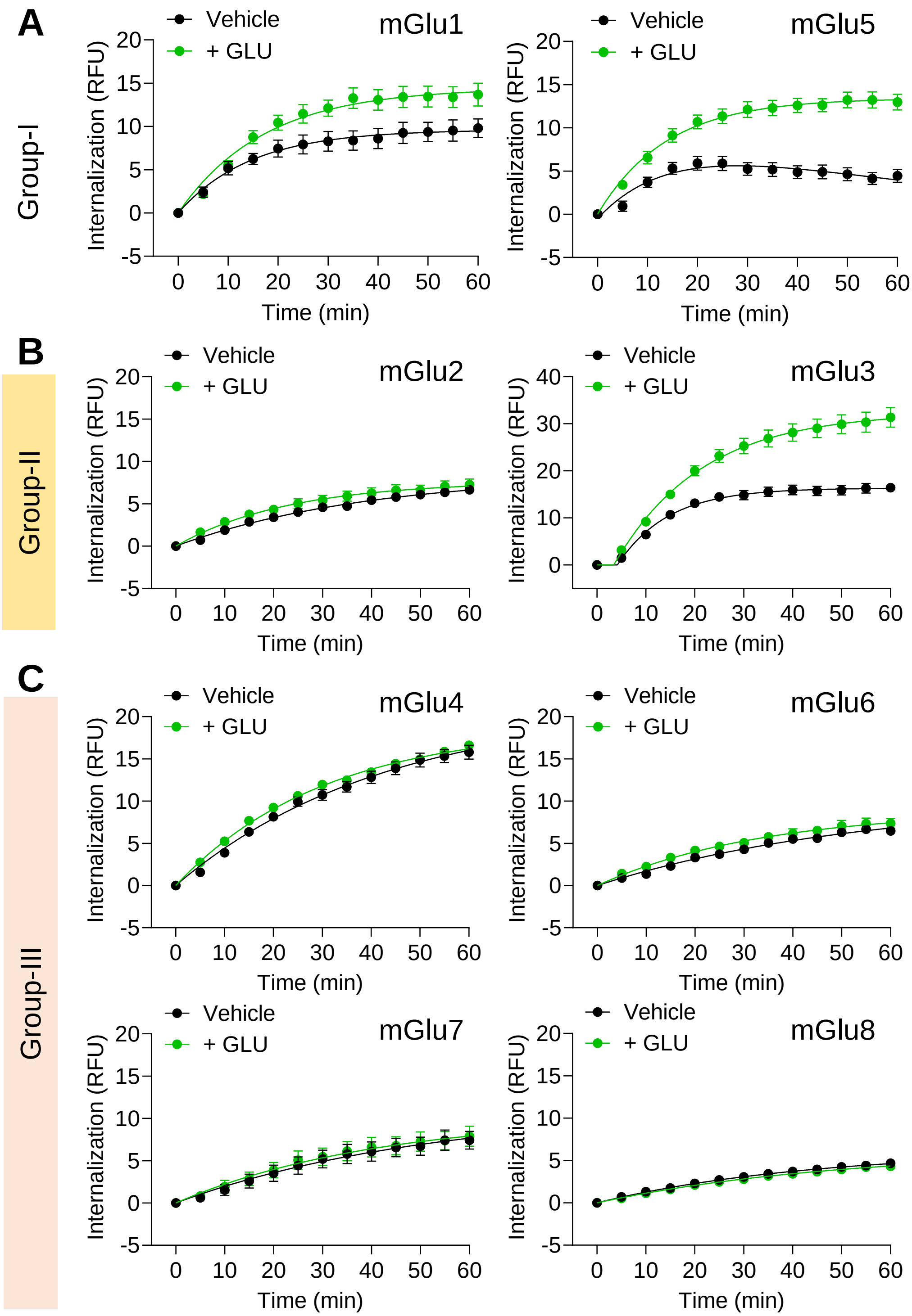


**Figure S4.** Internalization kinetics of mGlu receptors. (**A**) Internalization profiles of group-I mGlu receptors, mGlu1 and mGlu5 in the absence of agonist (vehicle) or 100 μM glutamate (+ GLU). (**B**) Internalization profile of group-II mGlu receptors, mGlu2 and mGlu3, in the absence of agonist (vehicle) or 100 μM glutamate (+ GLU). (**C**) Internalization profile of group-III mGlu receptors, mGlu4, mGlu6, mGlu7, mGlu8, in the absence of agonist in the absence of agonist (vehicle) or 100 μM glutamate (+ GLU). Data represent the mean ± S.E.M. of at least three independent experiments performed in triplicate.

### Fig S5

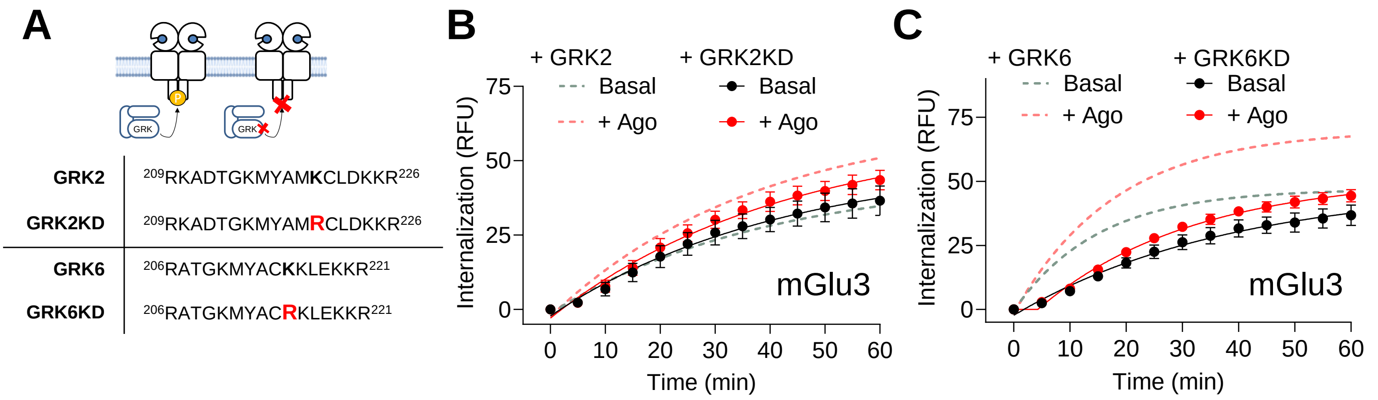


**Figure S5.** Kinase-dead GRKs impact on mGlu3 receptor internalization. (**A**) Schema of GRK-KD constructions: GRK2-K220R (GRK2-KD) and GRK6-K215R (GRK6-KD). (**B**) mGlu3 internalization kinetics in ΔGRK cells supplemented with GRK2 or GRK2-KD, in the absence of agonist (100 µM LY341495, Basal) or in the presence of 10 µM LY354740 (+ Ago). (**C**) mGlu3 internalization kinetics in ΔGRK cells supplemented with GRK6 or GRK6-KD, in the absence of agonist (100 µM LY341495, Basal) or in the presence of 10 µM LY354740 (+ Ago). Data represent the mean ± S.E.M. of at least three independent experiments performed in triplicate.

#### Fig S6

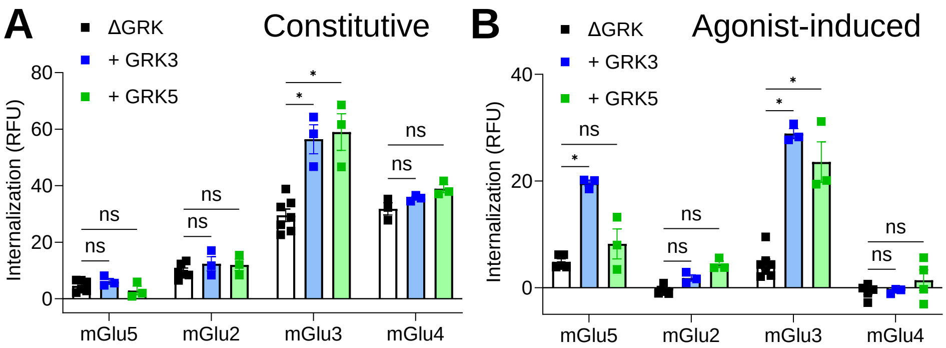


**Figure S6.** Effect of GRK3 and GRK6 on mGlu receptor internalization. (**A**) Constitutive internalization of mGlu5, mGlu2, mGlu3 and mGlu4 after 60 min of treatment with 100 μM LY341495, in ΔGRK cells (black), in ΔGRK cells + GRK3 (blue) or in ΔGRK cells + GRK5 (green). (**B**) Agonist-induced internalization of mGlu5, mGlu2, mGlu3 and mGlu4 after 60 min of treatment with a selective agonist (10 μM quisqualate for mGlu5, 10 μM LY354740 for mGlu2 and mGlu3, 10 μM LAP4 for mGlu4), in ΔGRK (black), in ΔGRK cells + GRK3 (blue) or in ΔGRK cells + GRK5 (green). Data represent the mean ± S.E.M. of at least three independent experiments performed in triplicate.

### Fig S7

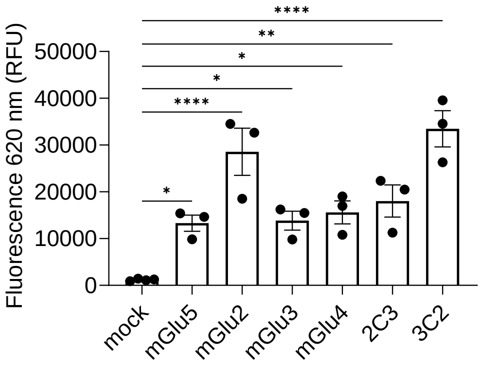


**Figure S7.** mGlu receptor expression measured after SNAP-lumi4-Tb labelling in Δβarr cells. Data represent the mean ± S.E.M. of at least three independent experiments performed in triplicate.

### Fig S8

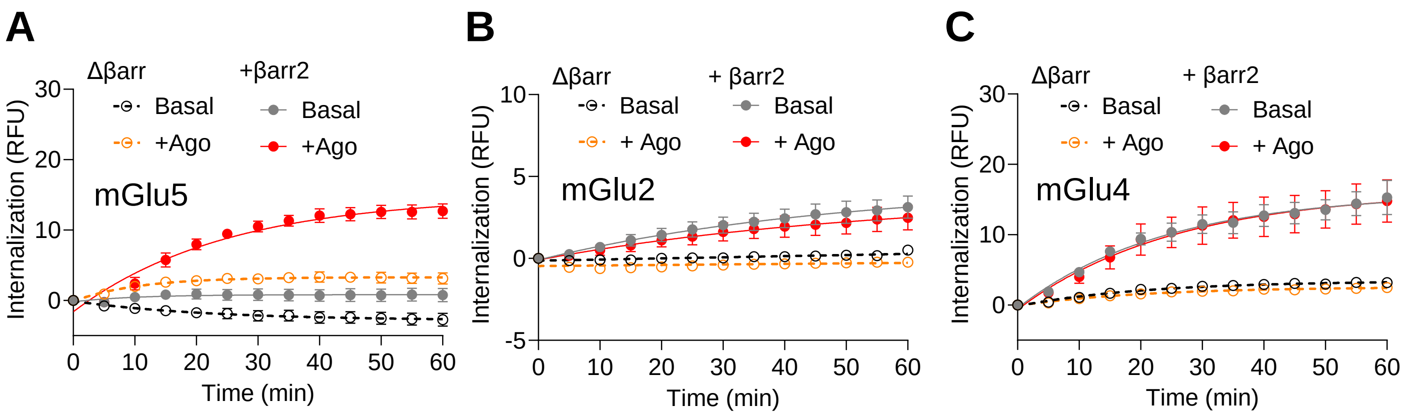


**Figure S8.** Internalization kinetics of mGlu receptors in cells lacking βarrs (Δβarr) and in Δβarr cells supplemented with βarr2 (+ βarr2). (**A**) mGlu5 internalization in the absence of agonist (100 μM LY341495, Basal) or 10 μM quisqualate (+ Ago). (**B**) mGlu2 internalization in the absence of agonist (100 μM LY341495, Basal) or 10 μM LY354740 (+ Ago). (**C**) mGlu4 internalization in the absence of agonist (100 μM LY341495, Basal) or 10 μM L-AP4 (+ Ago). Data represent the mean ± S.E.M. of at least three independent experiments performed in triplicate.

#### Fig S9

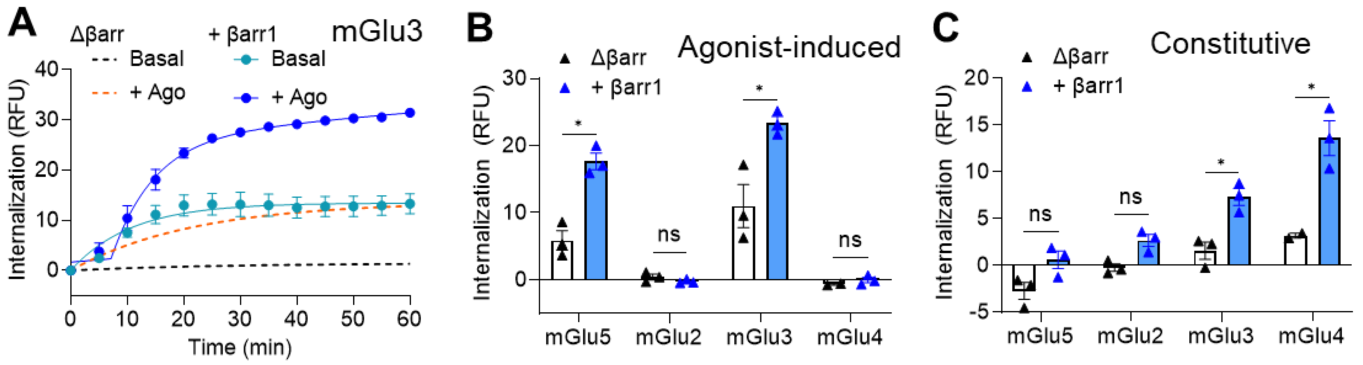


**Figure S9.** βarr1 effect on mGlu receptor internalization. (**A**) Internalization kinetics of mGlu3 receptor in Δβarr cells supplemented with βarr1 in the presence of 100 μM LY341495 (basal) or 10 μM LY354740 (+ Ago). (**B**) Agonist-induced internalization of mGlu5 (+ 10 μM quisqualate), mGlu2 (+ 10 μM LY354740), mGlu3 (+ 10 μM LY354740) or mGlu4 (+ 10 μM L-AP4) after 60 min in Δβarr cells (black bars) and in Δβarr cells supplemented with βarr1 (blue bars). (**C**) Constitutive internalization of mGlu5, mGlu2, mGlu3 and mGlu4 measured after 60 min of incubation with in 100 μM LY341495 in Δβarr cells (black bars) and in Δβarr cells supplemented with βarr1 (blue bars). Data represent the mean ± S.E.M. of at least three independent experiments performed in triplicate.

### Fig S10

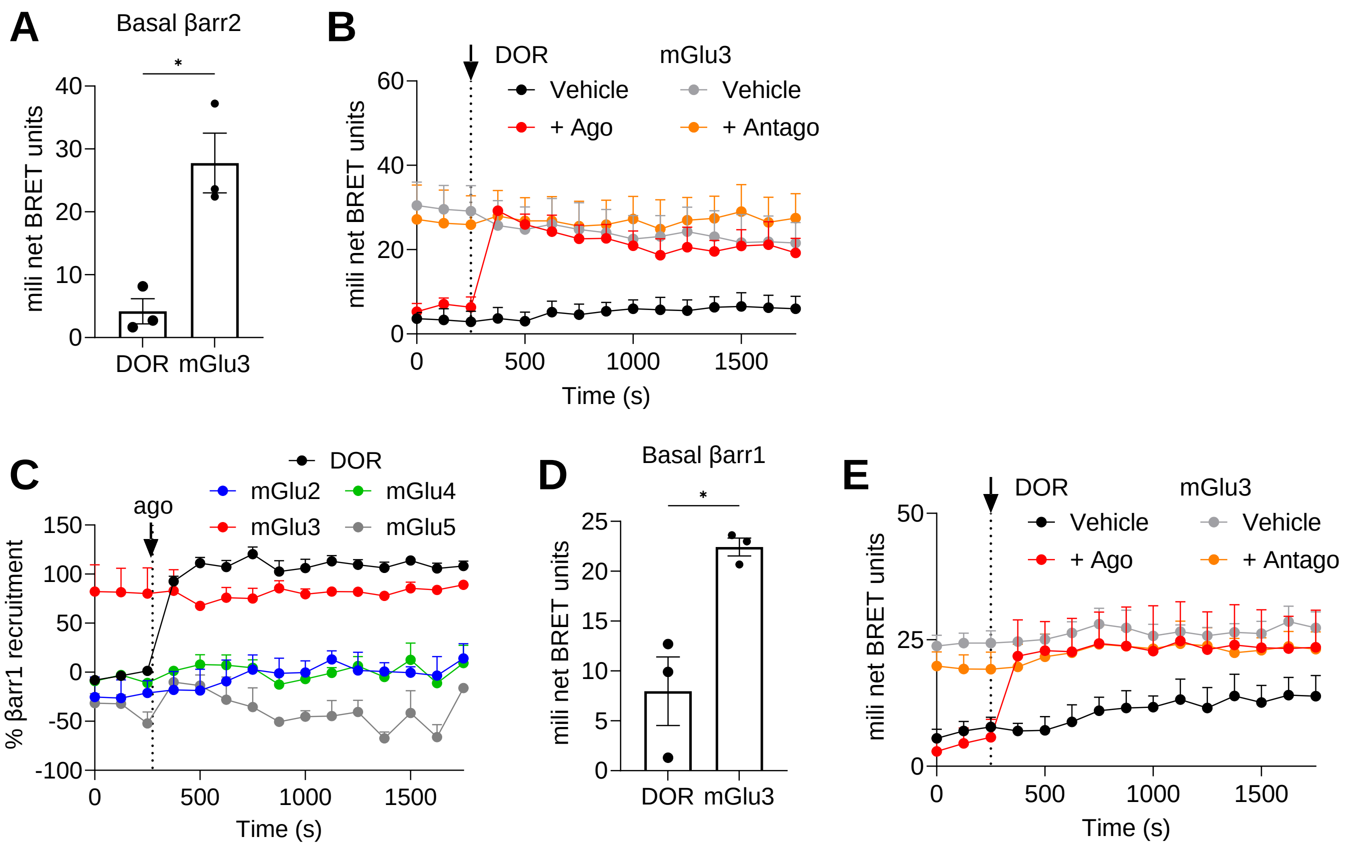


**Figure S10.** Control of BRET signal for βarr2 recruitment and effect of βarr1. **(A)** Basal BRET signal measured between βarr2-RLuc and mGlu3-Venus in the presence of 1 μM LY341495, compared with the basal signal obtained by DOR. (**B**) BRET kinetics of βarr2-RLuc recruitment in the absence of ligand (Vehicle) or after addition of 10 μM scn-162 (+ Agp) or 100 μM LY341495 (+ Antago). (**C**) mGlu-Venus recruitment of βarr1-RLuc measured by BRET before and after addition of a selective agonist (10 μM quisqualate for mGlu5, 10 μM LY354740 for mGlu2 and mGlu3, 10 μM LAP4 for mGlu4). Data are normalized by the maximal response obtained by the recruitment βarr2-RLuc by DOR-Venus in response to 10 μM snc-162. (**D**) Basal BRET signal measured between βarr1-RLuc and mGlu3-Venus in the presence of 1 μM LY341495, compared with the basal signal obtained by DOR. (**E**) BRET kinetics of βarr1-RLuc recruitment in the absence of ligand (Vehicle) or after addition of 10 μM scn-162 (+ Agp) or 100 μM LY341495 (+ Antago). Data represent the mean ± S.E.M. of three independent experiments performed in triplicate.

### Fig S11

**A**

| | |
|---|---|
| **(h)mGlu2** | HIILFQ$^{820}$**PQKNVVSHRAPTSRFGSAAARASSSLGQ GSGSQFVPTVCNGREVVDSTTSSL**$^{872}$ |
| **(h)mGlu3** | HIILFQ$^{829}$**PQKNVVTHRLHLNRFSVSGTGTTYSQSS ASTYVPTVCNGREVLDSTTSSL**$^{879}$ |
| **2C3** | HIILFQ$^{820}$**PQKNVVTHRLHLNRFSVSGTGTTYSQSS ASTYVPTVCNGREVLDSTTSSL**$^{870}$ |
| **3C2** | HIILFQ$^{829}$**PQKNVVSHRAPTSRFGSAAARASSSLGQ GSGSQFVPTVCNGREVVDSTTSSL**$^{881}$ |

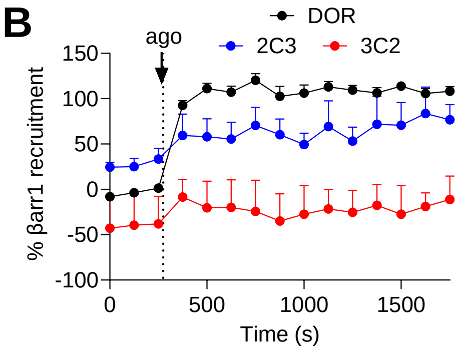


**Figure S11.** C-terminal exchange between mGlu2 and mGlu3. **(A)** Targeted sequences of mGlu2 and mGlu3 to build 2C3 and 3C2 constructs**. (B)** 2C3-Venus and 3C2-Venus recruitment of βarr1-RLuc measured by BRET before and after addition of a selective agonist (10 µM LY354740). Data are normalized by the maximal response obtained by the recruitment βarr2-RLuc by DOR-Venus in response to 10 µM snc-162. Data represent the mean ± S.E.M. of three independent experiments performed in triplicate.

### Fig S12

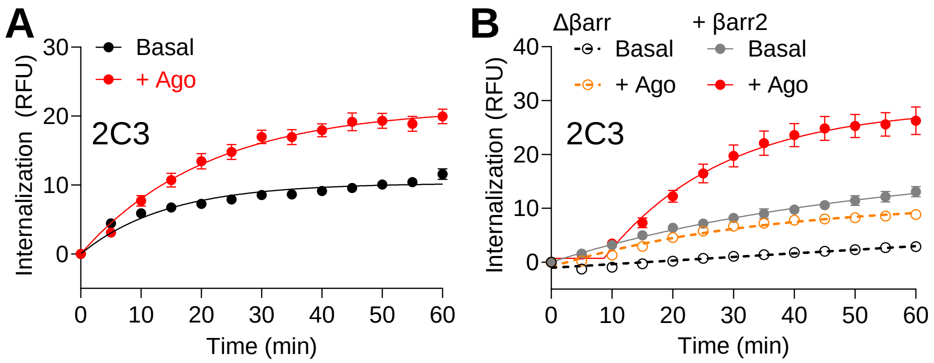


**Figure S12.** 2C3 internalization profiles. (**A**) Kinetic internalization of 2C3 in HEK293 cells in the absence of agonist (100 µM LY341495, Basal) or in the presence of 10 µM LY354740 (+ Ago). **(B**) Kinetic internalization of 2C3 in cells lacking βarrs (Δβarr) and in Δβarr cells supplemented with βarr2 (+ βarr2). Internalization was measured in the absence of agonist (100 µM LY341495, Basal) or in the presence of 10 µM LY354740 (+ Ago). Data represent the mean ± S.E.M. of three independent experiments performed in triplicate.

#### Fig S13

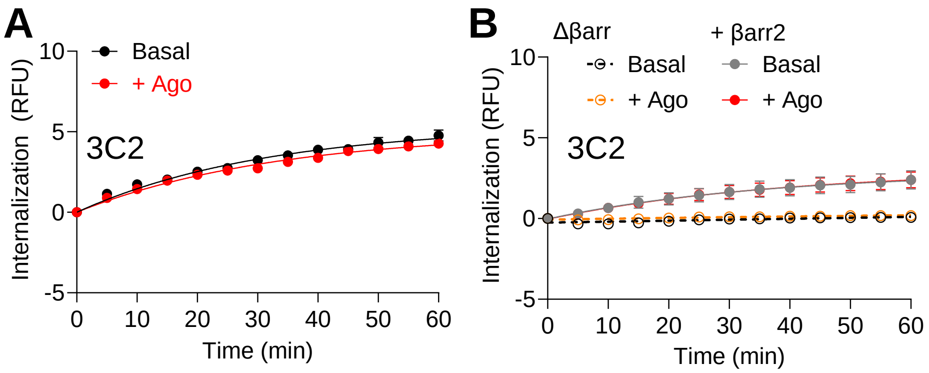


**Figure S13.** 3C2 internalization profile. (**A**) Kinetic internalization of 3C2 in HEK293 cells in the absence of agonist (100 μM LY341495, Basal) or in the presence of 10 μM LY354740 (+ Ago). **(B**) Kinetic internalization of 3C2 in cells lacking βarrs (Δβarr) and in Δβarr cells supplemented with βarr2 (+ βarr2). Internalization was measured in the absence of agonist (100 μM LY341495, Basal) or in the presence of 10 μM LY354740 (+ Ago). Data represent the mean ± S.E.M. of three independent experiments performed in triplicate.

### Fig S14

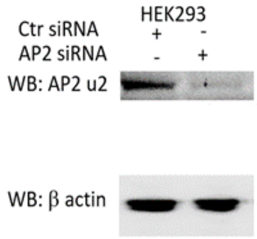


**Figure S14.** Detection of Western blot of the μ-subunit of AP2 in HEK293 cells treated with siRNA control Ctr siRNA) or siRNA-μAP2 by western blot. β-actin was included as loading control. Representative image out of three independent blots performed in triplicate.

### Fig S15

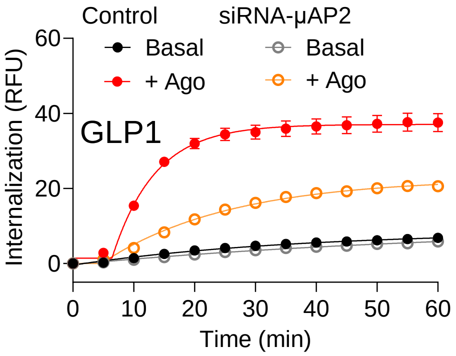


**Figure S15.** Internalization kinetics of GLP-1 measured in HEK293 cells (control) and HEK293 cells treated with siRNA-μAP2. Quantifications were done in the absence of agonist (Basal) or in the presence of 1 μM exendin-4 (+ Ago). Data represent the mean ± S.E.M. of three independent experiments performed in triplicate.

### Fig S16

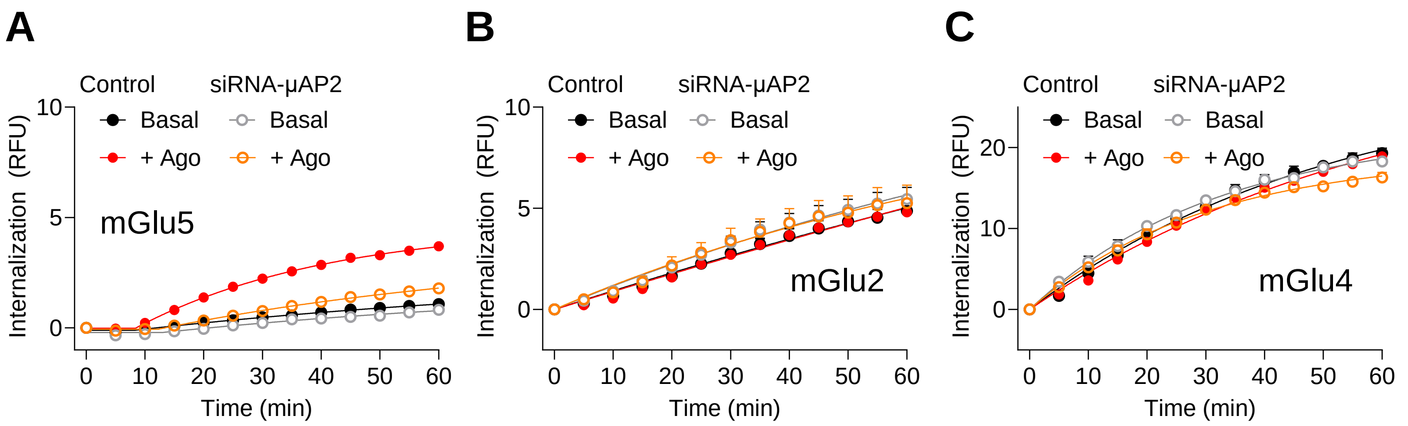


**Figure S16.** Downregulation of μAP2 subunit effect on internalization in control cells and cell with downregulated AP2**.** (**A**) mGlu5 internalization in the absence of agonist (100 μM LY341495, Basal) or 10 μM quisqualate (+ Ago). (**B**) mGlu2 internalization in the absence of agonist (100 μM LY341495, Basal) or 10 μM LY354740 (+ Ago). (**C**) mGlu4 internalization in the absence of agonist (100 μM LY341495, Basal) or 10 μM L-AP4 (+ Ago). Data represent the mean ± S.E.M. of at least three independent experiments performed in triplicate.

### Table S1

**Table S1**. Pharmacological parameters of internalization of mGlu5. Potency (pEC50) obtained after 60 min of internalization at 37 ºC. Data represent the mean ± S.E.M. of three independent experiments performed in triplicate.

| | QUIS | GLU | LY34 | LY34 (+ 0.1 μM QUIS) |
|---|---|---|---|---|
| $pEC_{50}$ | 7.64 ± 0.078 | 4.64 ± 0.20 | NA | 5.48 ± 0.13 |
